# Nonviral, ultrasound-triggered gene delivery platform via gas-core cationic nanobubbles

**DOI:** 10.64898/2026.09.11.749060

**Authors:** Laura E. Chen, Pinunta Nittayacharn, Aayushi Laliwala, Marcus Vincent Bella Jaro, Salima El Yakhlifi, Jean F. Eastman, Xu Han, Aaqib H. Khan, Kathryn HartMoore, Fatma Dogan, Valentina F. Giai, Timothy Brauns, Mark C. Poznansky, Richard K.P. Benninger, Marvin T. Nieman, Ilya Bederman, Mitchell L. Drumm, Agata A. Exner

## Abstract

Despite their promise, lipid nanoparticle gene delivery systems have repeatedly failed clinical trials and struggle to achieve efficient, localized transfection in target tissues. The majority of endocytosed nanoparticles are degraded before nucleic acid release, and an inability to track particle distribution *in vivo* prevents validation of successful delivery. Alternatively, nanobubbles (NBs) are lipid-shelled, gas-core preclinical ultrasound contrast agents and stimuli-responsive drug delivery vehicles. Under varying acoustic pressures, NBs expand, contract, and burst, releasing cargo in an externally controlled, site-specific manner while scattering unique echoes for simultaneous ultrasound visualization. Here, we introduce a cationic nanobubble (CNB) formulation with a +42.3 mV zeta potential, 265 nm diameter, and 2.43×10^11^ NBs/mL concentration. CNBs produce stable ultrasound contrast, electrostatically load plasmid DNA onto their surface, and internalize into >99% of human prostate cancer cells within 15 minutes *in vitro.* CNBs remain brightly echogenic intracellularly and induce sonication-dependent expression of green fluorescent protein (GFP). *In vivo*, CNBs generate contrast in mouse livers for 50 minutes after intravenous administration. Therapeutic ultrasound stimulation over the liver causes a sharp reduction in ultrasound contrast, visualizing localized cavitation in the target organ and inducing a 2.5-fold increase in anti-GFP mean fluorescence intensity relative to the untransfected control. Importantly, no GFP expression is observed without ultrasound stimulation, supporting a mechanism for selective and site-specific gene delivery. This study presents a highly stable CNB capable of efficient DNA loading and ultrasound-dependent gene expression. These results provide a foundation for the future development of CNB platforms to advance image-guided, ultrasound-triggered gene therapy.

## Introduction

Exogenous nucleic acid delivery alters cellular gene expression for a wide range of biomedical applications, serving both therapeutic and prophylactic roles across oncology, inherited disorders, and infectious disease^1^. Formulating nucleic acids into nanocarriers, such as liposomes and nanoparticles, is the predominant nonviral delivery strategy. While viral therapies are complicated by severe immune responses, mutagenesis of the host genome, and a costly production process, nanocarriers are less immunogenic and enable repeat dosing schemes and large-scale manufacturing^2–4^.

Although promising in preclinical models, nanocarriers have repeatedly failed human trials^5^. As of 2025, there are only four FDA-approved nanoparticle-based transfection agents for *in vivo* use: one for transthyretin-mediated amyloidosis treatment and three for viral immunization (SARS-CoV-2, respiratory syncytial virus)^2,6^. Several factors limit the translation of nanoparticle-mediated gene therapies, including low tissue accumulation, poor target specificity, and endosomal entrapment limiting effective transfection, as well as poor tracking of particle delivery *in vivo* preventing validation of successful delivery^7,8^.

Ultrasound is an emerging mechanical method to increase transfection efficiency. In addition to their applications in diagnostic imaging, acoustic waves can physically disrupt cell membranes, creating transient openings for drug or gene entry^9^. These mechanical effects can be further enhanced with the administration of ultrasound contrast agents: gas-core bubbles stabilized by a lipid or polymer shell. In response to varying acoustic pressures, shell-stabilized bubbles expand and contract in radius in the process of cavitation, nonlinearly scattering ultrasound waves to enhance image contrast^10^ and generating unique fluid streams and forces to open cellular junctions, permeabilize tissues, and trigger drug release^11^. Under mild acoustic stimulation, bubbles undergo stable cavitation and periodically oscillate in radius. At critical pressures, bubbles rapidly collapse inwards, sending microjets and shock waves into the surroundings via inertial cavitation. Ultrasound and bubble cavitation can also stimulate endocytosis, among several other bioeffects^9,12,13^.

Microbubbles (MBs) have been FDA-approved since the 1990s as diagnostic contrast agents, and MB sonoporation strategies are in development for gene delivery^14^. These ultrasound-mediated approaches include co-injection of MBs mixed with free nucleic acids, liposomes, or nanoparticles and loading genes directly onto to the MB itself^15,16^. However, due to their 1–10 µm diameter, MBs are largely restricted to the vasculature. As a result, MB-facilitated gene delivery poorly extends beyond endothelial cells, and transfection of deeper tissues often relies on the extravasation of co-delivered nanomaterials^17^.

Alternatively, submicron nanobubbles (NBs) are 100–600 nm diameter, preclinical contrast agents and drug delivery vehicles, whose small size permits bubble extravasation beyond endothelial barriers and increased cell uptake^18^. Leveraging NBs for gene delivery could provide several advantages over traditional lipid nanocarriers, including 1) increased tissue penetration and internalization^19,20^, 2) real-time, noninvasive imaging of particle distribution^21,22^, and 3) higher spatiotemporal control over delivery to minimize off-target effects^23^. Significantly, external cavitation of endocytosed NBs can also facilitate endosomal escape^24,25^. Together, these properties position ultrasound-stimulated NBs as unique tools for more effective, precise gene therapy.

NB gene delivery systems using positively-charged lipids, polymers, or chemical conjugation to load nucleic acids onto the bubble have been previously described^26^. However, many reports focus on therapeutic outcomes while deprioritizing a rigorous physicochemical characterization of the NB system. Even where preclinical therapeutic results are promising, practical clinical translation requires a detailed understanding of the NB gene carrier itself. Accordingly, in this work, we introduce a lipid-shelled, C_4_F_10_ gas-core cationic NB (CNB) formulation (**Fig. 1**), systematically assess particle surface charge, concentration, and size, and demonstrate the potential of CNBs as an ultrasound-mediated gene delivery platform. We characterize the CNBs with multiple orthogonal measurement techniques, including dynamic light scattering, resonant mass measurement, gas chromatography mass-spectroscopy, fluorescence microscopy, cryo-electron microscopy, and nonlinear contrast ultrasound imaging. To demonstrate proof-of-concept transfection, we load CNBs with plasmid DNA (pDNA) encoding green fluorescent protein (GFP) and evaluate cell uptake, echogenicity, and GFP expression in PC3 human prostate cancer cells *in vitro*. In addition, we assess *in vivo* ultrasound contrast longevity and transfection in mouse livers. Results demonstrate that CNBs enable nonviral, imageable gene delivery with ultrasound-triggered expression.

**Figure 1.**
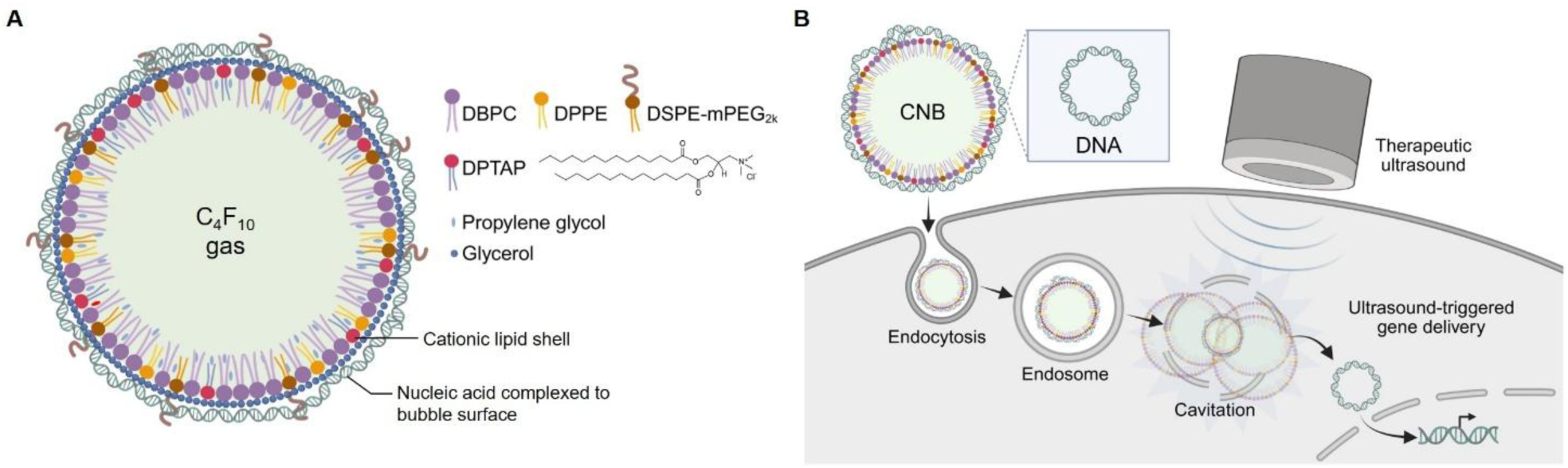
Cationic NBs (CNBs) for gene delivery. **A)** CNB formulation, consisting of a lipid shell and C_4_F_10_ gas core. Nucleic acids electrostatically load onto the cationic surface. **B)** Schematic of ultrasound-mediated gene delivery mechanism. Gene-loaded CNBs internalize into cells and cavitate under ultrasound stimulation to release the nucleic acid cargo.

## Results and Discussion

### CNB formulation and characterization

Our group has previously developed a stable, well-characterized NB formulation^27–29^ with several derivatives, including prostate-specific membrane antigen (PSMA)-targeted^30^ and chemotherapy-loaded^23^ contrast agents. These NBs have a net negative surface charge due to the phosphate head groups of the component lipids. In order to electrostatically load anionic nucleic acids, we first modified our NB to have a positive surface charge. The helper lipid 1,2-dipalmitoyl-sn-glycero-3-phosphate (DPPA) was replaced with the cationic lipid 1,2-dipalmitoyl-3-trimethylammonium-propane chloride (DPTAP), containing an equivalent 16-carbon saturated lipid tail but a positively charged trimethylammonium head group (**Fig. S1A**). The average zeta potential of DPTAP CNBs was 42.3 ± 1.9 mV (**Fig. 2A**) compared to −47.1 ± 2.8 mV of DPPA NBs (**Fig. S1B**), as measured in deionized water. By dynamic light scattering (DLS), CNBs were 267 ± 17 nm in number-weighted diameter and 279 ± 19 nm in intensity-weighted diameter (**Fig. 2B**) with a unimodal distribution (**Fig. 2C**) and polydispersity index (PDI) of 0.10 ± 0.02 (**Fig. 2D**). Particle size and concentration was also assessed by resonant mass measurement (RMM), which distinguishes buoyant, gas-core NBs from nonbuoyant particles such as micelles or lipoosomes^31^. By RMM, CNBs were 265 ± 17 nm in diameter (**Fig. 2E**) and 2.43 ± 0.92 ×10^11^ NBs/mL in concentration (**Fig. 2F**), comprising 95 ± 3.2% of total particles (**Fig. 2G**). Nonbuoyant particles were present with a 246 ± 43 nm average diameter at 9.58 ± 4.5 ×10^9^ particles/mL. To further confirm the presence of internalized gas, gas chromatography mass-spectroscopy (GC/MS) was also performed^32^. C_4_F_10_ characteristically splits into CF_3_^+^, C_2_F_7_^+^, and C_3_F_5_^+^ molecules upon ionization. The area under the curve (AUC) of CF_3_^+^ ion abundance versus time increased linearly with the number of CNBs (**Fig. 2H, I**). To visually confirm size, CNBs were flash-frozen and imaged via cryo-electron microscopy (cryo-EM). Circular, lipid-shelled structures were observed with diameters consistent with the distributions found by DLS and RMM (**Fig. 2J**). For fluorescence tracking, rhodamine-CNBs (rCNBs) were also produced using a rhodamine-conjugated lipid by the same methods. rCNBs had similar physical properties to the non-fluorescent CNBs, with an average zeta potential of 40.3 ± 2.2 mV (**Fig. S2A**), number-weighted diameter of 278 ± 11 nm, intensity-weighted diameter of 296 ± 18 nm (**Fig. S2B, C**), and PDI of 0.11 ± 0.05 (**Fig. S2D**) by DLS. By RMM, rCNBs were 255 ± 33 nm in diameter at 1.23 ± 0.17 ×10^11^ NBs/mL (**Fig. S2E, F**) with a 97 ± 2.5% buoyancy (**Fig. S2G**).

**Figure 2.**
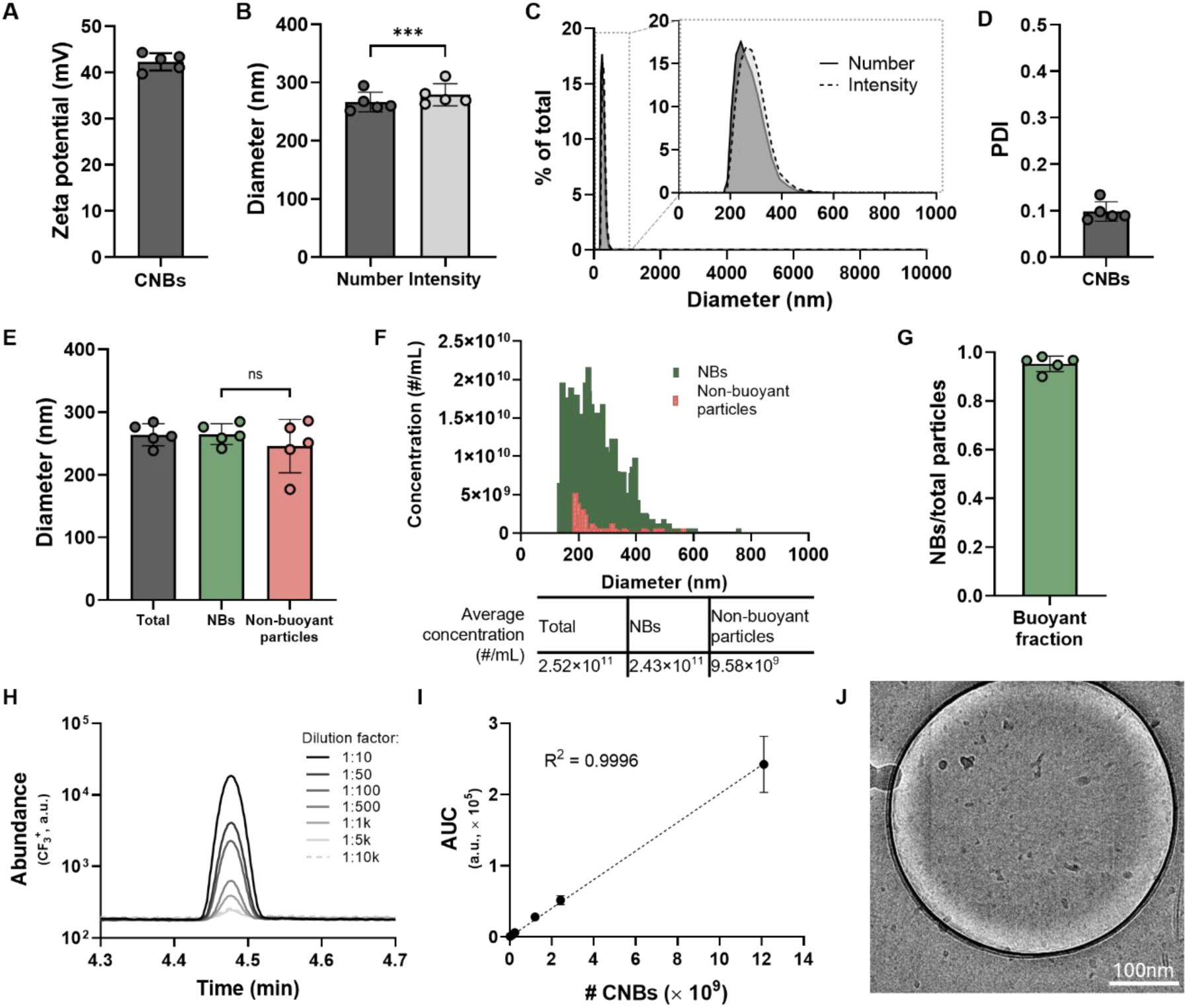
Characterization of cationic nanobubbles (CNBs). **A)** Zeta potential in DEPC-treated water (n = 5). **B)** Peak number-weighted and intensity-weighted diameters by dynamic light scattering (DLS) (n = 5). Significance by two-tailed paired t test, ***p < 0.001. **C)** Average percent size distribution by DLS with inlay highlighting 0–1000 nm range (n = 5). **D)** Polydispersity index (PDI) by DLS (n = 5). **E)** Average diameter of buoyant NBs, non-buoyant particles, and total particles by resonant mass measurement (RMM) (n = 5). **F)** Concentration vs. diameter distributions of NBs and non-buoyant particles by RMM (n = 5). **G)** Fraction of buoyant NBs to total particles by RMM (n = 5). **H)** Gas chromatography mass-spectroscopy plots of CF_3_^+^ (*m*/*z* = 69) ion abundance versus time at various CNB dilutions with **I)** corresponding area-under-the-curve (AUC) quantification vs. number of CNBs, R^2^ = 0.9996 (n = 3). **J)** Cryo-electron microscopy of a CNB. Scale bar 100 nm.

Lipid structure is crucial to NB stability, ultrasound responsiveness^33^ and payload delivery^34^. While other lipid NB formulations have used DC-cholesterol^35,36^ or DOTAP^37^ as the cationic component, DPTAP was selected for highest structural compatibility with our preestablished formulation and produced highly concentrated, C_4_F_10_-core bubbles with a uniform, nanoscale size distribution. DPTAP^38^ and analogs^16^ have also been used in MBs, and the lipid maintains a permanent positive charge at physiological pH^39^. Alternatively, lipid nanoparticles commonly use ionizable lipids for transfection, which transition from neutral to positively charged in acidic environments to promote endosomal escape^7,40^. While we considered incorporating ionizable lipids into the CNB, their often branched, unsaturated tail structures could destabilize the CNB shell and weaken acoustic properties. Additionally, our gene-delivery system proposes an active, externally triggered cargo release with ultrasound. This allows for site-specific transfection instead of solely relying on the passive endosomal escape of ionizable lipids. Furthermore, given the NB gas core limits internal cargo, nucleic acids are loaded onto the surface; thus, a permanent cationic charge is favorable to keep the cargo electrostatically complexed while in circulation. In future studies, other cationic lipid candidates and combinations will be explored to further modulate NB properties and gene release.

### CNB pDNA loading capacity by gel electrophoresis

Having produced nanoscale bubbles with a cationic charge, we next evaluated their ability to load pDNA. To capture the ratio between loaded pDNA mass and CNB number, we expressed pDNA concentration in micrograms of pDNA per milliliter of undiluted CNBs, denoted µg/NBmL. This notation ensures the ratio of pDNA to CNBs is kept consistent, even when CNBs may be diluted differently in phosphate-buffered saline (PBS) depending on experimental needs. CNBs were incubated with a 3,486 base-pair plasmid encoding green fluorescent protein (pmax-GFP) at 0–1000 µg/NBmL, then samples were run on an agarose gel. pDNA-CNB complexes will be retained in the sample well, while free pDNA will migrate toward the positive electrode. Signal enhancement of the sample wells increased with pDNA concentration, as did free DNA band intensity (**Fig. 3A**). When band intensity was quantified against a standard curve (**Fig. S3**), >90% of total pDNA was bound to the CNBs at a 250 µg/NBmL loading concentration and below, while 60 ± 9% was bound at 500 µg/NBmL and 47 ± 13% of pDNA was bound at 1000 µg/NBmL (**Fig. 3B**). This corresponds to a maximum total CNB loading capacity of 469 ± 125 µg/NBmL (**Fig. 3C**), or approximately 2 fg of DNA per individual CNB.

**Figure 3.**
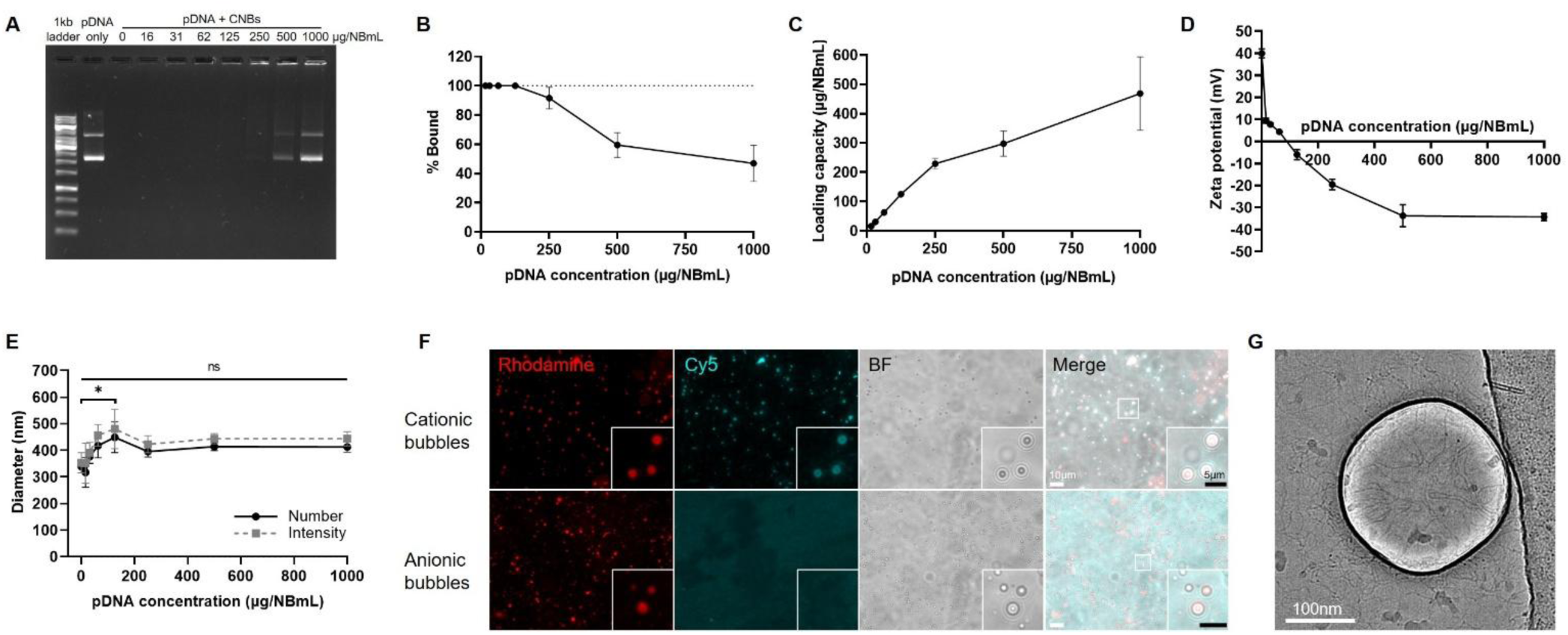
Plasmid DNA loading onto CNBs. CNBs were incubated with a range of pmax-GFP pDNA concentrations (expressed as µg of pDNA per mL of undiluted NBs, µg/NBmL) spanning 0–1000 µg/NBmL and run on an **A)** agarose gel for quantification of **B)** percent bound DNA (n = 3) and **C)** DNA loading capacity (n = 3). **D)** Zeta potential vs. pDNA concentration (n = 3). **E)** Peak number-weighted and intensity-weighted diameters by DLS vs. pDNA concentration (n = 3). Significance by one-way ANOVA with multiple comparisons relative to the unloaded control. \**p* < 0.025. **F)** Fluorescence microscopy of Cy5-pDNA complexed to rhodamine cationic bubbles compared to anionic bubbles. Microbubbles instead of NBs were imaged to accommodate the resolution limits of the microscope. Scale bar 10 µm. Inlay scale bar 5 µm. **G)** Cryo-electron microscopy of a CNB loaded with 250 µg/NBmL pDNA. Scale bar 100 nm.

By qualitative visual inspection of agarose gels, other CNB designs using DC-cholesterol have reported a loading capacity of 1 µg plasmid to 15–20 µL of their NBs^41,42^. While exact NB concentration was not reported in these studies, this would be equivalent to a loading capacity of 50-67 µg/NBmL in our notation. Therefore, in relation to other lipid-based CNBs, the DPTAP formulation could provide a 7–9-fold higher pDNA loading capacity.

### Size and charge characterization of pDNA-loaded CNBs

CNB zeta potential became more negative as pDNA loading concentration increased, transitioning from cationic to anionic between 62.5 µg/NBmL and 125 µg/NBmL before plateauing to a minimum charge of −34.3 ± 1.7 mV when incubated at 1000 µg/NBmL (**Fig. 3D**). The zeta potential plateau was consistent with the loading capacity trend by gel electrophoresis and suggests successful plasmid complexation to the CNB surface. An increase in CNB diameter was observed between 0–125 µg/NBmL, followed by a slight decrease at 250 µg/NBmL and subsequent plateau (**Fig. 3E**). Differences in diameter were not statistically significant, except between the unloaded and 125 µg/NBmL condition (number-weighted *p* = 0.024; intensity-weighted *p* = 0.011). An increase in particle diameter is expected as more pDNA is complexed to the surface; it is possible that free pDNA in the sample or changes in electrostatic repulsion caused a lower size estimation by DLS for the highest pDNA conditions.

### Fluorescence and cryo-electron microscopy confirm pDNA complexation to CNB surface

To visually confirm that pDNA electrostatically localized to the bubble surface, a Cy5 fluorophore was covalently conjugated to the plasmid before loading onto cationic versus anionic rhodamine bubbles. Bubbles were not size-isolated prior to pDNA loading in this experiment to capture the larger MBs resolvable by fluorescence microscopy. Cy5-pDNA formed around the periphery of cationic bubbles but not anionic bubbles (**Fig. 3F**). pDNA-CNBs were also imaged by cryo-EM. Strands of DNA accumulated around the edges and surface of the CNBs (**Fig. 3G**). Some DNA was also observed in the surrounding ice, which could represent free plasmid or displaced plasmid from the cryo-EM grid glow discharging process.

Unlike most nanoparticles, CNBs have an internal gas core and utilize external loading of DNA onto their surface. This loading technique leaves the cargo partially exposed to the external environment, where nucleases could degrade vulnerable nucleic acids^43^. Surface-loading of pDNA onto other cationic MB^44–46^ and NB^44^ formulations has been shown to protect against DNase degradation, though this concept remains to be tested with the CNBs presented here. If required, several methods could be explored to improve protection of the cargo, such as coating with cross-linked polymers^47^ or nesting the pDNA-CNB within a secondary lipid shell^48^.

### CNBs produce stable ultrasound contrast *in vitro*

After investigating the CNBs’ physical properties and pDNA loading capacity, we next evaluated their *in vitro* acoustic response in an agarose, tissue-mimicking phantom at 18 MHz, 4% power, and 1 frame-per-second (fps) (**Fig. S4**). CNBs produced bright nonlinear contrast in a concentration-dependent manner (**Fig. 4A**), and pDNA loading did not significantly affect initial signal intensity (**Fig. 4B**). During 500 seconds of continuous exposure to the imaging transducer, the unloaded (**Fig. 4C**) and pDNA-loaded (**Fig. 4D**) CNBs maintained constant signal intensity over time when diluted to 10^9^ NBs/mL in PBS. Signal decayed to 82 ± 8.3% of the original at 10^8^ NBs/mL and 55 ± 19% of the original at 10^7^ NBs/mL for unloaded CNBs. For pDNA-loaded CNBs, signal intensity after 500 seconds was 76 ± 12% of the original at 10^8^ NBs/mL and 31 ± 17% of the original at 10^7^ NBs/mL.

**Figure 4.**
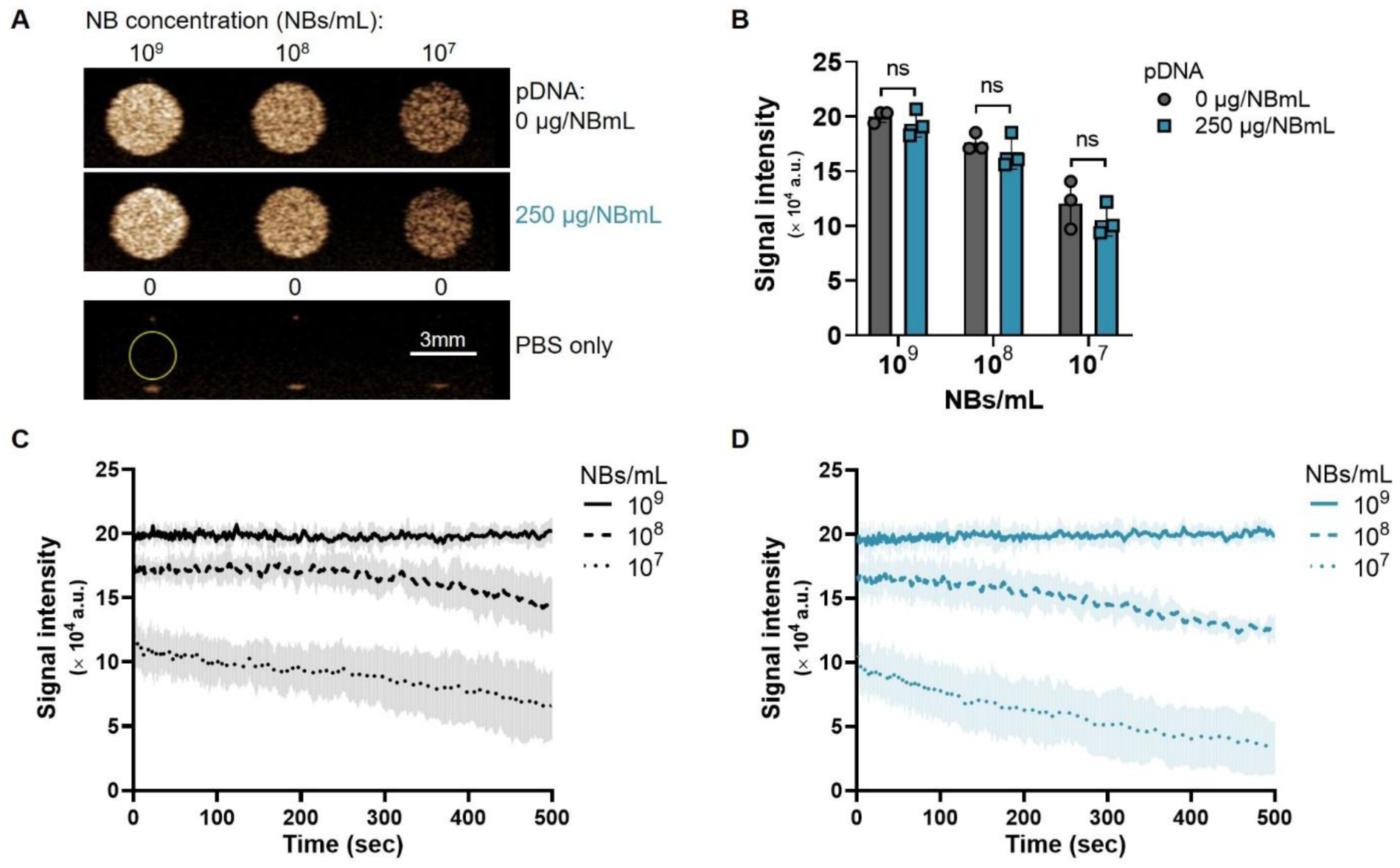
Ultrasound contrast characteristics of unloaded and pDNA-loaded CNBs *in vitro*. **A)** Representative nonlinear contrast images of CNBs loaded with 0 µg/NBmL or 250 µg/NBmL of pDNA (µg of pDNA per mL of undiluted NBs) after dilution to the order of 10^9^, 10^8^, or 10^7^ NBs/mL in PBS. PBS-only wells and representative region of interest shown. Corresponding quantification of **B)** initial signal intensity and **C,D)** signal intensity over time for **C)** unloaded CNBs and **D)** pDNA-loaded CNBs (n = 3). PBS background subtracted.

To assess longitudinal stability, nonlinear contrast intensity was periodically assessed over 144 hours (6 days) post NB isolation under storage at 4°C and ambient pressure. Both CNBs (**Fig. S5A, C**) and rCNBs (**Fig. S5B, D**) maintained visible contrast over time. After 144 hours, signal intensity decreased to 57 ± 20% of the original for CNBs and 63 ± 12% of the original for rCNBs when measured at 10^9^ NBs/mL (**Fig. S5E**).

The rate of ultrasound signal decay is an indirect measurement of bubble stability; as gas diffuses out of the NB core over time, echogenicity is reduced. These results demonstrate the CNBs are stable and produce high contrast intensity under continuous ultrasound stimulation, after several days of storage at 4°C, and when loaded with pDNA. NB shell composition, stiffness, and cargo loading is known to influence oscillatory response and stability in an acoustic field^27,49^. Plasmid loading did not significantly affect contrast intensity in these experiments; however, future passive cavitation detection studies could more precisely evaluate changes in acoustic response^50^. Storing the CNBs in a pressurized, C_4_F_10_ gas-saturated vial instead of at ambient pressure could further extend their longevity, and other preservation techniques such as freeze-drying could also be investigated^51^.

### Unloaded rCNBs rapidly internalize into PC3 cells *in vitro*

To assess the cellular uptake and cytotoxicity of the CNBs, adherent PC3 human prostate cancer cells were incubated with 5,000, 10,000, or 20,000 rCNBs/cell over 180 minutes and analyzed by flow cytometry. PC3 cells were used given our group’s prior experience characterizing theranostic, anionic NBs in this cell line^25,30,52^. rCNBs had no negative effect on cell viability over a 3-hour incubation (**Fig. 5A**). Within 5 minutes, >97% of cells were rhodamine positive when exposed to 5,000 NBs/cell, and >99% of cells were rhodamine positive when exposed to 10,000 or 20,000 NBs/cell (**Fig. 5B**). Median fluorescence intensity (MFI) of the rhodamine positive cells increased over time and followed a NB concentration-dependent trend (**Fig 5C**). Cell populations uniformly increased in fluorescence (**Fig. 5D**), and uptake was also confirmed by microscopy (**Fig. 5E**).

**Figure 5.**
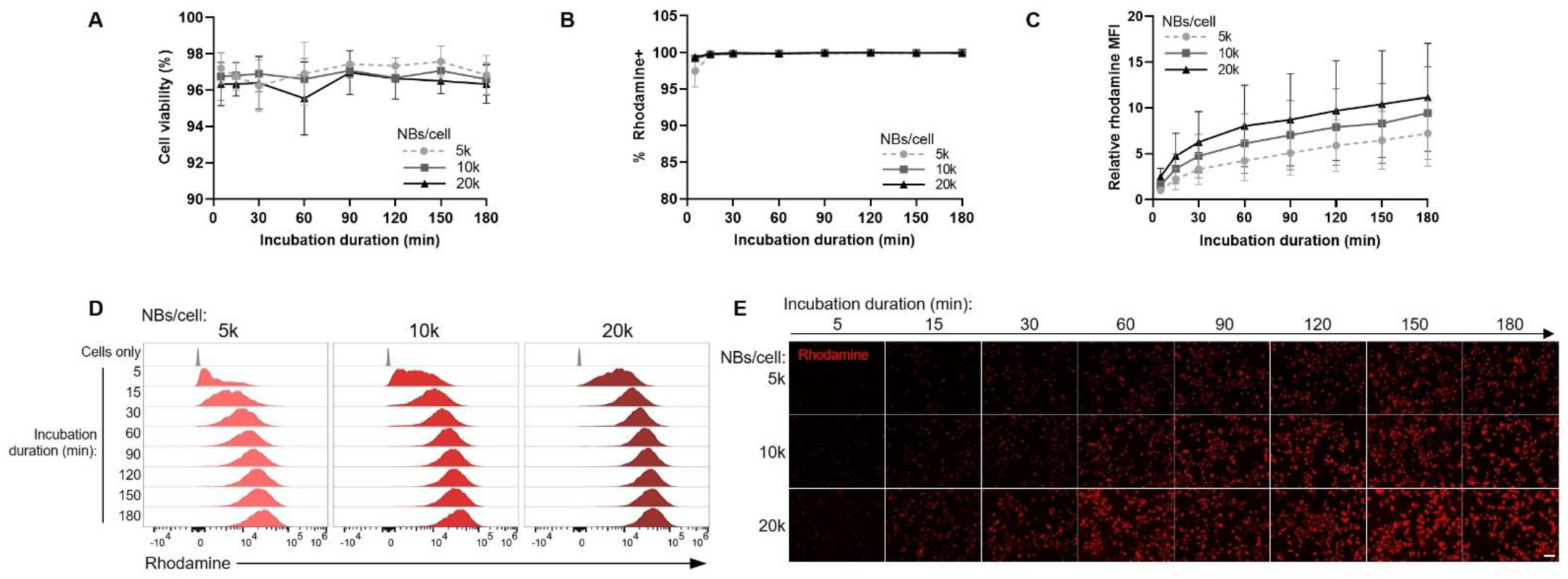
Uptake of unloaded rhodamine-CNBs in PC3 prostate cancer cells *in vitro*. Cells were incubated with 5k, 10k, or 20k rCNBs/cell over 180 minutes. Flow cytometry quantification (n = 3) of **A)** percent cell viability, **B)** percent rhodamine positive cells, **C)** median fluorescence intensity (MFI) of rhodamine positive cells as a fraction of the 5k NBs/cell, 15-minute internal control, and **D)** cell distribution histograms of rhodamine fluorescence. **E)** Representative rhodamine fluorescence microscopy images of PC3 cells over time. Scale bar 100 µm.

Nanomaterial internalization is governed by many factors, such as particle size, charge, stiffness, shape, hydrophobicity, and component chemistry^53,54^. Transfection agents favor cationic lipids’ electrostatic interaction with phospholipid cell membranes; however cationic compounds can also induce harmful reactive oxygen species and cytosolic leakage^55,56^. Favorably, our rCNBs internalized into >99% of cells within 15 minutes and demonstrated no baseline cytotoxicity *in vitro* over the time period evaluated. As shown later in Fig. 8C, no negative impact on cell viability was also observed after a 24-hour exposure.

### pDNA-loaded rCNBs internalize into cells in a surface charge-dependent manner

After demonstrating strong internalization of cargo-free rCNBs, we then assessed PC3 cell uptake kinetics of the rCNBs when loaded with 50, 125, or 250 µg/NBmL of pDNA. Similar to the 0 µg/NBmL condition, pDNA-rCNBs had no negative impact on cell viability (**Fig. 6A**). The lowest 50 µg/NBmL loading condition experienced the most rapid cellular uptake, internalizing into >99% of cells after a 15-minute incubation (**Fig. 6B**) and producing a higher MFI than the 125 and 250 µg/NBmL conditions across all timepoints (**Fig. 6C–E**). pDNA loading inhibited cell uptake in a concentration-dependent manner. Averaged across all timepoints, the rhodamine MFI of cells incubated with pDNA-CNBs at 50 µg/NBmL was 86% of the unloaded control, and the 125 and 250 µg/NBmL pDNA-CNBs produced an MFI of 38% and 30% of the unloaded control, respectively (**Fig. 6D**). Reduced cell uptake with higher pDNA loading is expected, as CNB zeta potential decreases and will interact less readily with negatively charged cell membranes. To maximize CNB uptake, we proceeded with the 50 µg/NBmL loading concentration for subsequent transfection experiments.

**Figure 6.**
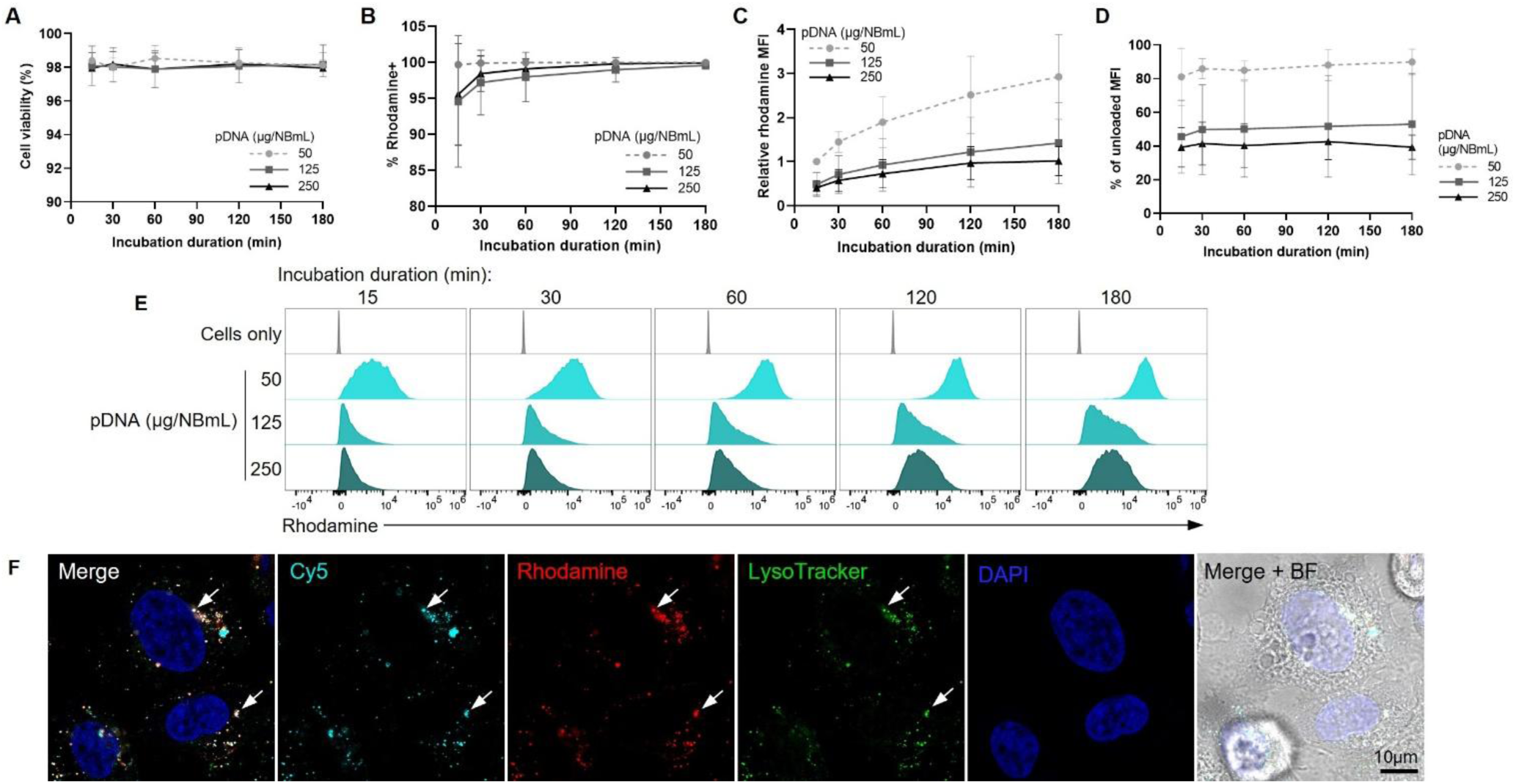
Uptake of pDNA-loaded rhodamine-CNBs in PC3 prostate cancer cells *in vitro*. Cells were incubated with 20k rCNBs/cell loaded with 50, 125, or 250 µg/NBmL pDNA (µg of pDNA per mL of undiluted NBs) over 180 minutes. Flow cytometry quantification (n = 3) of **A)** percent cell viability, **B)** percent rhodamine positive cells, **C)** median fluorescence intensity (MFI) of rhodamine positive cells as a fraction of the 50 µg/NBmL, 15-minute internal control, **D)** MFI as a percentage of the 0 µg/NBmL MFI at each timepoint, and **E)** representative cell distribution histograms of rhodamine fluorescence. **F)** Confocal microscopy of PC3 cells after a 15-minute incubation with rCNBs loaded with 50 µg/NBmL of Cy5-pDNA, including LysoTracker stain for late endosomes/lysosomes and DAPI nuclear stain. Arrows point to examples of Cy5, rhodamine, and LysoTracker colocalization. Scale bar 10 µm.

### CNBs deliver pDNA intracellularly

The flow cytometry uptake experiments used rCNB fluorescence as a proxy for pDNA internalization. To directly confirm rCNBs were facilitating pDNA entry, pDNA was also labeled with a Cy5 fluorophore prior to rCNB loading at 50 µg/NBmL, and cells were imaged by confocal microscopy with a LysoTracker stain for late endosomes/lysosomes. Distinct regions of rCNB, Cy5-pDNA, and LysoTracker colocalization were observed throughout the cell cytoplasm (**Fig. 6F**). Instances of Cy-5 pDNA that did not overlap with rCNBs or LysoTracker were also observed, suggesting some pDNA could have dissociated from the rCNBs or been internalized separately. These results demonstrate that the CNBs facilitate intracellular pDNA entry *in vitro* without any ultrasound intervention.

### PC3 cells produce nonlinear contrast after CNB internalization

To support the flow cytometry results of CNB uptake and confirm cell-associated CNBs were ultrasound-responsive, the nonlinear contrast intensity of PC3 cells after CNB incubation was evaluated in an agarose phantom. Cells with no CNBs were visible on B mode imaging but not in nonlinear contrast mode (**Fig. 7A**). After a 15-minute incubation with CNBs, cells displayed bright nonlinear contrast that increased with the number of CNBs per cell. Signal intensity of the cell suspension after a 1-hour exposure to CNBs was lower than a 15-minute exposure (**Fig. 7A,B**). To assess the longevity of intracellular CNB signal, excess CNBs were washed off after 15 minutes, and cells were left for an additional hour at 37°C before ultrasound imaging. Average signal intensity decreased >75% over the 1-hour incubation. To confirm that ultrasound could cavitate intracellular CNBs, CNB-laden PC3 cells were also exposed to 3 MHz therapeutic ultrasound in suspension at varying intensities and durations before imaging (**Fig. 7C, S6**). Nonlinear contrast intensity of the cells decreased with increasing ultrasound exposures (**Fig. 7D**). Approximately 50% of the initial contrast remained after the highest 2.2 W/cm^2^, 60-second exposure.

**Figure 7.**
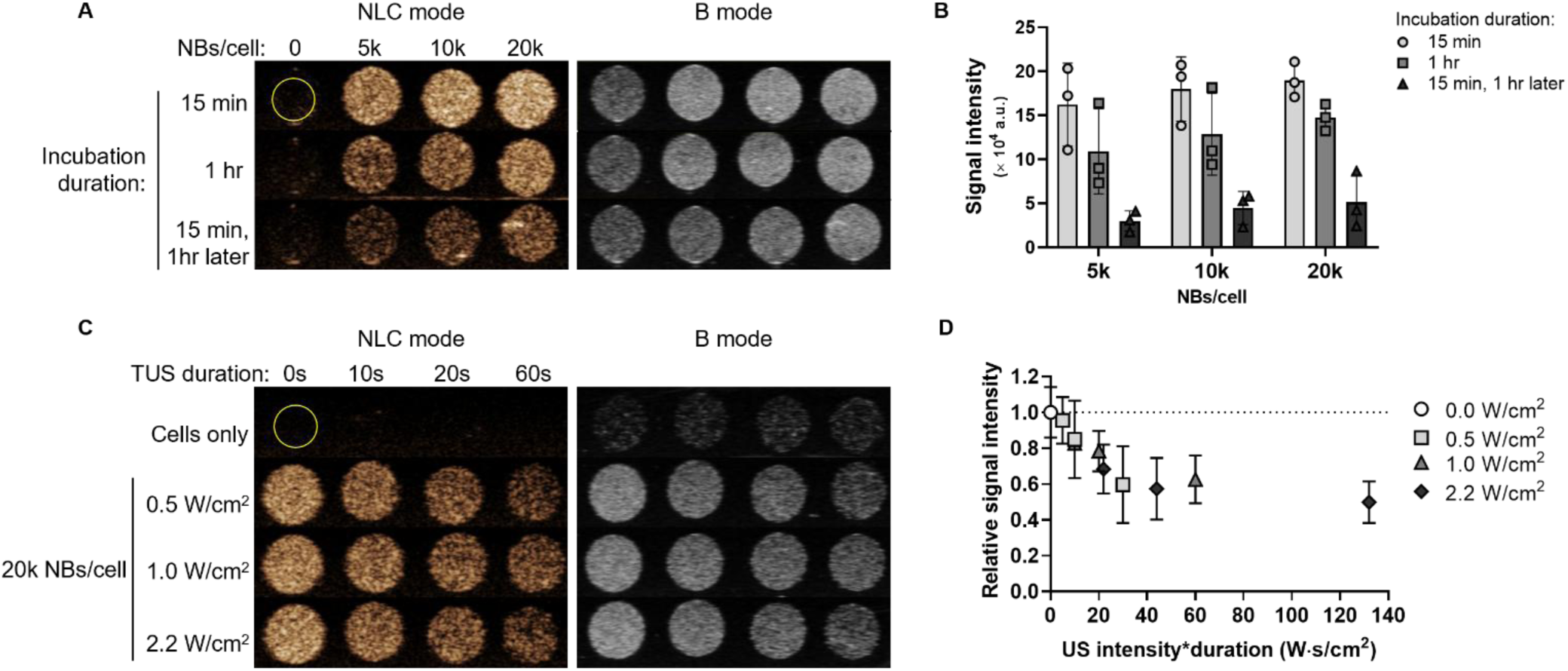
Ultrasound imaging of PC3 cells after CNB internalization and cavitation. **A)** Representative nonlinear contrast (NLC) and B-mode images of PC3 cells in suspension at 1×10^6^ cells/mL after incubation with 5k, 10k, or 20k unloaded CNBs/cell at 37°C for 15 minutes, 1 hour, or 15 minutes with 1 additional hour in incomplete RPMI with **B)** quantification of nonlinear contrast intensity (n = 3). Cells-only background subtracted. **C)** Representative NLC and B-mode images of PC3 cells at 100k cells/mL after a 15-minute incubation with 20k NBs/cell and therapeutic ultrasound (TUS) exposure for 0, 10, 20, or 60 seconds at 0.5, 1.0, or 2.2 W/cm^2^ with **D)** quantification of nonlinear contrast intensity versus ultrasound intensity*duration relative to the no TUS (0.0 W/cm^2^) control (n = 3). Cells-only background subtracted.

While particle uptake monotonically increased with time according to rhodamine fluorescence, cells were more echogenic after a 15-minute incubation with CNBs compared to a 1-hour incubation. This finding is likely due to gas diffusion out of the CNBs over time, decreasing their response by nonlinear contrast imaging but not affecting their fluorescence detection by flow cytometry. Therapeutic ultrasound exposure to the cells decreased signal intensity, indicating CNB cavitation occurred. Further passive cavitation studies could classify the presence of stable versus inertial cavitation; however, the low ultrasound intensities used (calculated pressure equivalents of 121 kPa, 172 kPa, and 255 kPa) would suggest stable cavitation^50^. Additionally, cells retained nonlinear contrast after sonication, indicating the persistence of intact CNBs.

Optimal transfection will require balancing maximum particle uptake before loss of the NB gas core to maintain ultrasound responsiveness. Previous work by our group has demonstrated that NBs targeted to PSMA can persist within intracellular vesicles for over 24 hours, maintaining detectable echogenicity^52^. While the PSMA-targeted NBs have a negative surface charge and are primarily internalized by receptor-mediated endocytosis, these CNBs achieved high uptake without any receptor-specific binding. To extend the intracellular longevity of the CNBs, adding a targeting receptor could be explored in future studies.

### Ultrasound-dependent GFP expression *in vitro*

After determining pDNA-CNBs successfully internalized into PC3 cells and remained echogenic, we next assessed the ability of an ultrasound trigger to induce expression of GFP *in vitro* (**Fig. 8A**). GFP expression was evident by microscopy 24 hours post-transfection for ultrasound-exposed groups; however, no GFP expression was observed in cells that received pDNA-rCNBs with no ultrasound (**Fig. 8B, S7**). By flow cytometry, cell viability mildly decreased with increasing ultrasound intensity*duration to a minimum of 87 ± 6.4% at the highest exposure (**Fig. 8C**). Over 99% of cells were rhodamine positive across all groups (**Fig. 8D**). Transfection efficiency was calculated as the percentage of GFP positive live cells out of the total number of live cells. Without ultrasound stimulation, just 0.13 ± 0.03% of live cells were GFP positive (**Fig. 8E**). With ultrasound, transfection efficiency peaked at 4.0 ± 2.6% (range 1.8–6.8%) after 10 seconds of exposure at 2.2 W/cm^2^. Relative GFP MFI also highest for the 2.2 W/cm^2^, 10-second condition (**Fig. 8F**). To investigate the role of internal endosome cavitation versus external membrane perforation in facilitating DNA expression, a comparison experiment with anionic NBs was also performed. Anionic NBs will not electrostatically complex with pDNA and will internalize less-readily compared to cationic particles; therefore, GFP expression in this condition is expected to primarily result from membrane permeabilization rather than endosomal cavitation. When mixed with an equivalent amount of pDNA and cavitated at 2.2 W/cm^2^ for 10 seconds, anionic NBs produced a transfection efficiency of 0.83% ± 0.65% (**Fig. S8**).

**Figure 8.**
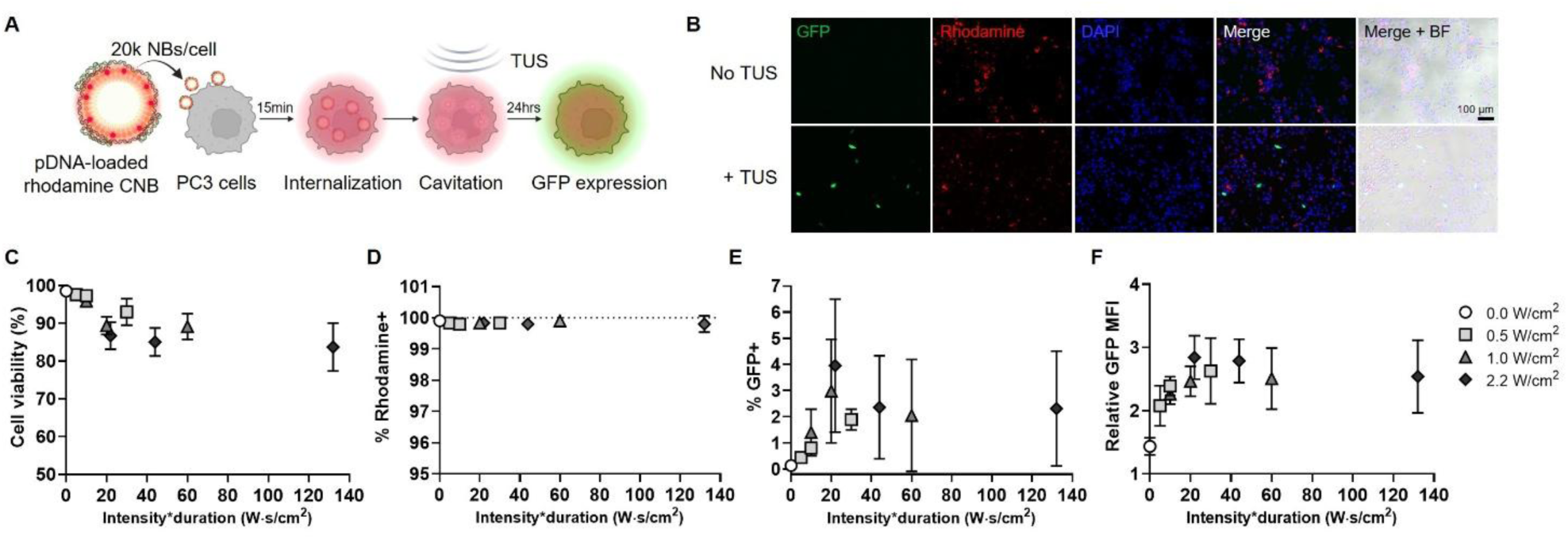
*In vitro* transfection of PC3 prostate cancer cells with pmax-GFP pDNA-loaded rCNBs. **A)** Schematic of transfection protocol. **B)** Representative fluorescence microscopy images 24 hours post-transfection with and without therapeutic ultrasound (TUS) stimulation at 3 MHz, 20% DC, 0.5 W/cm^2^ for 60 seconds (see Fig. S7 for complete microscopy at all TUS parameters). Scale bar 100 µm. Flow cytometry quantification of **C)** cell viability, **D)** percent rhodamine positive cells, **E)** percent GFP positive cells, and **F)** relative GFP median fluorescence intensity (MFI) of live, GFP positive cells vs. ultrasound intensity*duration.

There are several methods of assessing transfection efficiency, including observational microscopy^44,57^ and flow cytometry^58^ after fluorescent reporter expression, luminescence after luciferase expression^38,45,59–61^ and fold-copy number reduction after siRNA knockdown^37,62^. It is challenging to compare transfection efficiencies across *in vitro* studies, as they have analysis- and cargo-specific functional outcomes and use drastically different amounts of nucleic acid. Comparably, MBs have achieved *in vitro* expression of fluorescent reporter proteins in the 0.2–30% range^17,63,64^. While our uptake studies suggested >99% of cells internalized the pDNA-rCNBs, this did not translate to the same fraction of cells expressing GFP. There are several possible reasons for this finding. First, the *in vitro* suspension environment will cause variation in ultrasound exposure across individual cells, and not all CNBs could have been sufficiently cavitated with the parameters used. Secondly, even if released into the cytosol, DNA must also cross the nuclear membrane for mRNA transcription. Studies into DNA uptake after MB-enhanced sonoporation have suggested ∼30% of internalized DNA reaches the nucleus, while the remainder is left in the cytoplasm or autophaogosomes^65^. Nuclear entry is generally highest during mitosis, when the dissolved nuclear envelope reforms during telophase^66^; therefore, if cell proliferation was impaired by ultrasound exposure, this could also lead to lower DNA expression. Future studies delivering mRNA instead of DNA could investigate the degree of cytoplasmic gene delivery without the nuclear membrane barrier. Additionally, we observed a decrease in cell number with increasing ultrasound exposures, likely from cell lysis and dissolution. The peak in GFP expression at an intermediate ultrasound intensity*duration suggests a trade-off between achieving sufficient CNB cavitation for gene release and preserved cell integrity. One proposed mechanism for this gene release is CNB cavitation within endosomes, supported by the high degree of pDNA-CNB uptake before ultrasound intervention and the comparably low transfection efficiency with anionic NBs. Ultimately, the *in vitro* cell suspension does not mimic the physiological environment where gene-loaded NBs would act. To utilize NBs’ unique deformability, tissue penetration, and real-time imaging properties, we proceeded with characterization *in vivo*.

### CNBs produce lasting contrast *in vivo* that locally diminishes after ultrasound cavitation

To assess CNB stability under physiological conditions, nonlinear contrast in mouse livers was imaged over time after intravenous administration. CNBs produced immediate, high signal intensity that persisted for over 50 minutes above baseline under continuous imaging (**Fig. 9A, B**). On average, pDNA-CNBs produced slightly higher contrast than unloaded CNBs; however, this difference was not statistically significant (*p* = 0.52 at 50 minutes). After 5 minutes of local therapeutic ultrasound over the liver, a sharp reduction in nonlinear contrast was observed in the sonicated region (**Fig. 9C, D**). No acute toxicity reactions were observed after CNB administration, and all mice recovered fully from the procedure.

**Figure 9.**
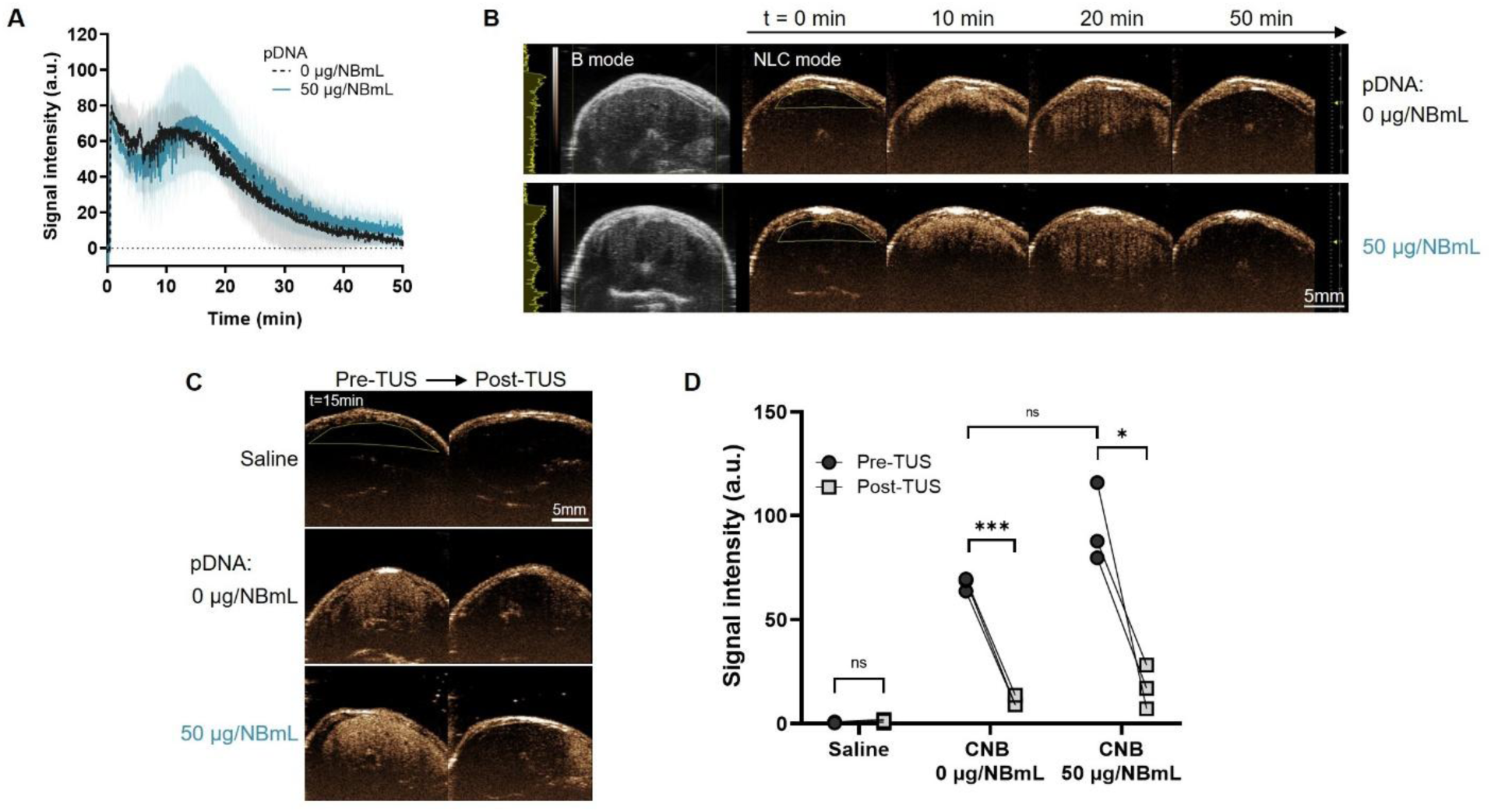
*In vivo* ultrasound imaging of CNBs in mice. **A)** Nonlinear contrast intensity in the liver over time after a 200 µL bolus intravenous injection of CNBs diluted 2-fold in PBS and loaded with 0 µg/NBmL or 50 µg/NBmL pDNA (µg of pDNA per mL of undiluted NBs) (average n = 3, standard deviation shaded). Pre-injection background subtracted. **B)** Representative B-mode and nonlinear contrast (NLC) images, with quantified region of interest shown. Scale bar 5 mm. **C)** Nonlinear contrast images of the liver 15 minutes post-injection and immediately after therapeutic ultrasound (TUS) application at 3 MHz, 2.2 W/cm^2^, 20% DC for 5 minutes. Scale bar 5 mm. **D)** Quantification of liver nonlinear contrast intensity immediately before and after TUS (n = 3). Significance between pre- and post-TUS groups by two-tailed paired t-test, \*\*\**p* < 0.001, \**p* < 0.05. Between-group significance by two-tailed unpaired t-test.

The *in vivo* environment introduces many factors that could affect bubble performance, including higher temperature^67–69^, dilution into the intravascular volume^28,70^, and interaction with plasma proteins and red blood cells^71^. Generally, cationic particles have a low circulation time due to rapid protein opsonization and immune cell phagocytosis, and electrostatic disruption of cell membranes is a concern for biocompatibility^72–74^. Notably, this CNB formulation produced no acute reactions in mice and provided long-lasting contrast in the liver for 50 minutes under continuous imaging. The pDNA-CNBs had a lower cationic charge compared to the unloaded CNBs, which could explain to their slightly higher contrast over time. The reduction in nonlinear contrast after therapeutic ultrasound suggested successful, localized cavitation of the CNBs over the target organ.

### Ultrasound-dependent GFP transfection in mouse livers

After nonlinear contrast imaging provided an immediate indication of gene release, mice were kept for 24 hours to allow time for GFP expression (**Fig. 10A**). The combination of pDNA-rCNBs and therapeutic ultrasound induced a 2.5 ± 0.49 -fold increase in anti-GFP AF647 MFI in the liver compared to the saline control (*p* = 0.002) by confocal microscopy (**Fig. 10B, D**). Relative AF647 MFI after pDNA-CNB administration was 2.3-fold higher with therapeutic ultrasound compared to without (*p* = 0.003). Without therapeutic ultrasound, pDNA-rCNBs produced no significant GFP expression (*p* = 0.99). Unloaded rCNBs produced a 1.7 ± 0.21 -fold increase in anti-GFP AF647 MFI (*p* = 0.11) after cavitation. To detect rCNB distribution, rhodamine fluorescence was also assessed in the liver, and a few punctate remnants were observed (**Fig. 10C, D**).

**Figure 10.**
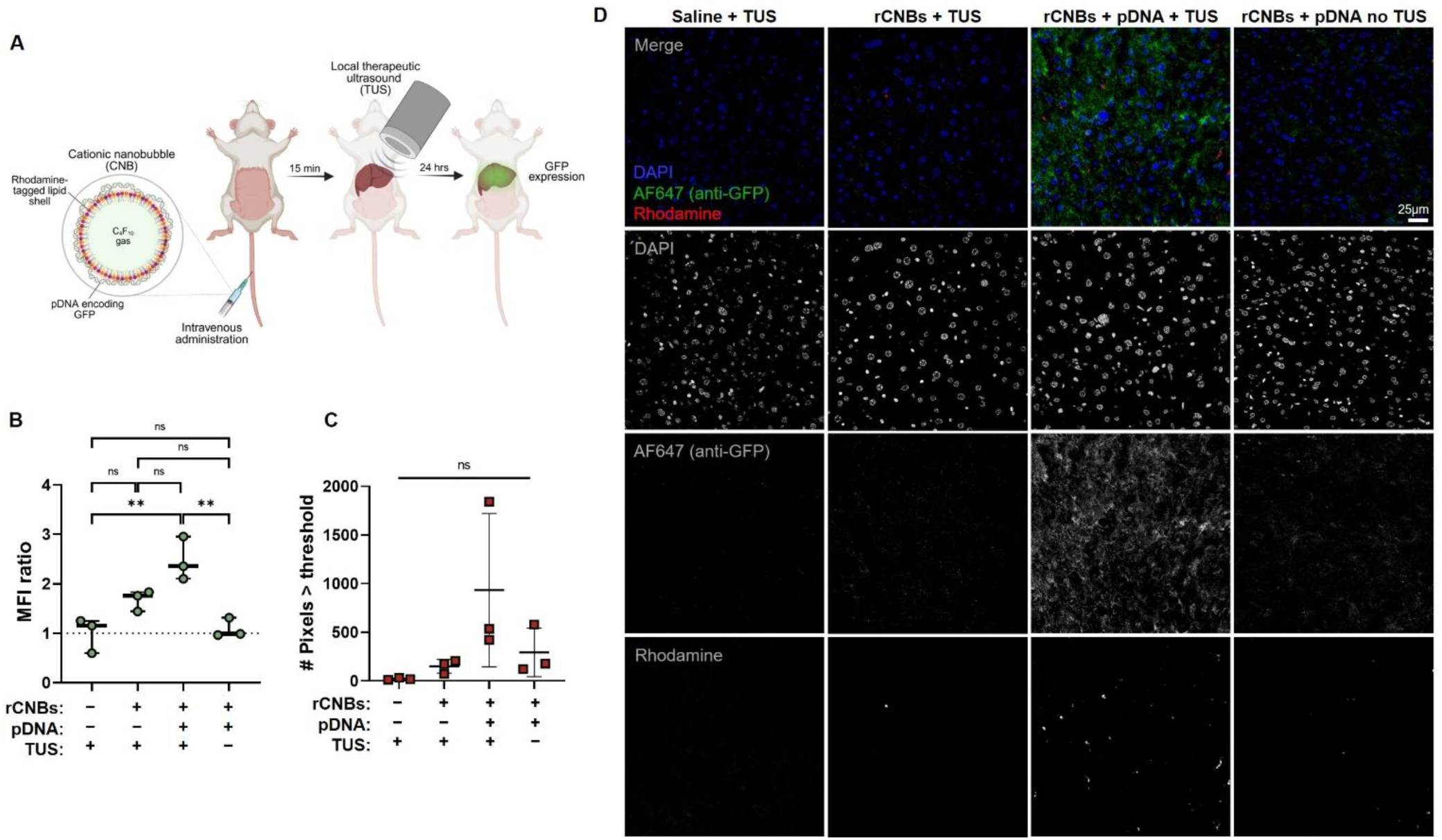
*In vivo* rCNB transfection in mouse livers. **A)** Schematic of transfection protocol. Quantification of **B)** average anti-GFP AF647 mean fluorescence intensity (MFI) and **C)** average number of rhodamine pixels above threshold in liver slices by confocal microscopy, 24 hours post-transfection with rCNBs loaded with 0 µg/NBmL or 50 µg/NBmL pmax-GFP pDNA, with or without therapeutic ultrasound (TUS). MFI is normalized to cell count in the image and expressed as a ratio to the saline-TUS control (n = 10 fields-of-view for each mouse, n = 3 mice per group). Significance by one-way ANOVA with multiple comparisons. \*\**p* < 0.005. **D)** Representative fluorescence confocal microscopy images of livers, including DAPI (nuclei), AF647 (anti-GFP), and rhodamine (rCNBs). Scale bar 25 µm.

These results demonstrate ultrasound-dependent transfection with pDNA-loaded CNBs *in vivo*. No GFP expression occurred without ultrasound stimulation, which could prevent off-target transfection in future applications. The liver was selected as the target organ for proof-of-concept transfection in this study; however, it is important to note that many nanocarriers are naturally sequestered in the liver through protein adsorption and uptake by the reticuloendothelial system^75^. Future studies will explore transfection in tumor models and extrahepatic organs, which historically have been the more challenging targets for gene delivery platforms^76^. A statistically insignificant but interesting increase in MFI was observed with unloaded rCNBs and therapeutic ultrasound. This could be the result of higher metabolic activity or stress in the tissue, as oxidation products can demonstrate high autofluorescence^77^. Given no MFI increase was observed in the pDNA-rCNB group without ultrasound, pDNA loading could be important to help shield the cationic charges. The low rhodamine signal suggests most of the rCNB lipid shells had been cleared from the liver after 24 hours.

## Conclusions

This study presents a highly stable CNB for ultrasound-triggered gene delivery. Robust characterization of particle size, charge, nucleic acid loading capacity, and echogenicity lays a supportive groundwork for future experimentation and therapeutic translation. CNBs generate bright, long-lasting nonlinear contrast within cells *in vitro* and after intravenous administration *in vivo*. Ultrasound imaging offers a real-time, noninvasive method to monitor particle distribution, and an immediate reduction in contrast after therapeutic ultrasound is consistent with localized CNB cavitation in the target organ. Despite nearly all prostate cancer cells internalizing the pDNA-CNBs *in vitro*, no significant GFP expression was observed without ultrasound stimulation. Although additional mechanistic studies are required, these findings support a model in which cavitation of intracellular pDNA-CNBs promotes cytoplasmic gene delivery. GFP expression was also ultrasound-dependent in mouse livers *in vivo,* suggesting the potential for spatially controlled gene expression with precise sonication of the target. While transfection of reporter DNA was demonstrated here, further work is necessary to evaluate the safety, mechanism, and efficacy of these CNBs in disease models. By swapping the nucleic acid cargo and moving the ultrasound transducer over the tissue of interest, we hope this platform could be adapted to address multiple clinical needs.

## Methods

### NB production

CNBs were produced with the lipids 1,2-dibehenoyl-sn-glycero-3-phosphocholine (DBPC) (Avanti),1,2-dipalmitoyl-sn-glycero-3-phosphoethanolamine (DPPE) (Avanti), 1,2-distearoyl-sn-glycero-3-phosphoethanolamine-N-[methoxy(polyethylene glycol)-2000] (DPSE-mPEG_2k_) (LaysanBio), and 1,2-dipalmitoyl-3-trimethylammonium-propane chloride (DPTAP) (Avanti) in a 6:2:1:1 mass ratio (**Fig. S1A**). A lipid emulsion was formed in propylene glycol and heated to 82 °C for 1 hour. A PBS-glycerol solution was then added to achieve a final lipid concentration of 1 mg/mL. To form bubbles, 1 mL of the lipid emulsion was placed in a sealed vial, and air was exchanged with C_4_F_10_ gas. The vial was mechanically agitated for 45 seconds (VialMix) to create a polydisperse bubble population, and bubbles of nanometer-scale were isolated through differential centrifugation at 50 g for 5 minutes.

Anionic NBs were produced in accordance with previous methods^27,29^ with an equivalent mass of the anionic 1,2-dipalmitoyl-sn-glycero-3-phosphate (DPPA) instead of cationic DPTAP. To produce rhodamine fluorescent bubbles, 50 μg/mL of 1,2-dipalmitoyl-sn-glycero-3-phosphoethanolamine-N-(lissamine rhodamine B sulfonyl) (DPPE-rhodamine) (Avanti) was supplemented to the lipid mixture before production.

### Characterization of NB size, charge, and concentration

CNB size distribution and PDI were measured by DLS (Anton Paar Litesizer 500). NBs were diluted to the order of 10^8^ NBs/mL in PBS and read in a 1.5 mL polystyrene cuvette (Fisher Scientific) at 25°C. Zeta potential (Anton Parr Litesizer 500) was measured at 10^8^ NBs/mL in deionized DEPC-treated water (Ambion Invitrogen) in an omega cuvette (Anton Paar) at 25°C. Water was used as the solvent instead of PBS to prevent excess ions in solution from obscuring the surface charge. CNB concentration was measured via RMM (Archimedes, Malvern Panalytical)^31^ with a MicroH-A6303H sensor. NB positively-buoyant particle density was set to 0.008 g/mL, and negative buoyancy was set at 1.34 g/mL. CNBs were diluted 10,000-fold in PBS and measured until a minimum of 500 particles were collected. The average CNB concentration of n = 5 replicates was used for subsequent NB-to-cell ratio calculations.

### Gas chromatography mass-spectroscopy

To confirm the presence of encapsulated C_4_F_10_ gas, CNBs were diluted in PBS (10, 50, 100, 500, 1000, 5000, and 10000-fold) to 500 μL total volume in GC/MS vials. To release gas into the headspace, sealed vials were sonicated for 20 minutes in a 50 °C water bath. Samples were analyzed by a GC/MS platform (Agilent Technologies) consisting of a 5977B mass spectrometer coupled with a 7890B gas chromatograph. Briefly, 1 μL of the gas headspace was injected into the gas chromatograph at 1:20 split ratio and 200°C injector temperature, and analytes were separated using a HP-5MS capillary column (30 m × 250 μm × 0.25 μm) maintained under 1.5 mL/min helium flow. The oven temperature program was as follows: 60°C held for 1 minute, then ramp of 40°C/min until 120°C, then held for an additional 3.5 minutes. The detector was held at 250°C. The most abundant fragment, CF_3_^+^ (*m/z* = 69), was quantified at a retention time of 4.48 minutes. Additional characteristic fragments of C_4_F_10_, including C_2_F_5_⁺ (*m/z* = 119) and C_3_F_7_⁺ (*m/z* = 169), were also detected. All fragment ions were confirmed using the NIST mass spectral database.

### Cryo-EM sample preparation and imaging

To prepare samples for cryo-EM, Quantifoil R2/2 holey carbon grids (300 mesh) were glow-discharged using an EMITech K100X Glow Discharge unit for 60 seconds at 25 mA and 0.2 mBar. All samples were frozen using an FEI Mark IV Vitrobot maintained at 4°C and 100% humidity. A 2.5 μL aliquot of CNBs was applied to the grid and allowed to incubate for 30 seconds. Grids were then blotted with standard Vitrobot filter paper (Ted Pella) for 10 seconds at a blot force of 5. After blotting, samples were immediately vitrified in liquid ethane, then stored under liquid nitrogen until cryo-EM imaging. The Titan Krios G3i Cryo-TEM operating at 300 kV equipped with a K3 direct electron detector was used to collect data in super-resolution mode at 64,000 magnification (physical pixel size: 1.34 Å/pix, 0.66 Å/pix for super-resolution, total dose: 50 e^-^/Å^2^). Data were collected using Serial EM and motion corrected in the cryoSPARC V4.2.1 suite.

### Plasmid DNA production

Plasmid DNA encoding green fluorescent protein (Lonza pmax-GFP, 3,486 base pairs) with kanamycin-resistance and a cytomegalovirus promoter was purified with the GeneJET Plasmid MaxiPrep Kit (ThermoFisher Scientific) according to manufacturer’s protocol. DNA was stored at −20°C and freeze-thaw cycles were minimized to maintain plasmid integrity.

### Plasmid DNA loading onto CNBs

pmax-GFP pDNA was loaded onto the CNBs by electrostatic association. By our convention, the loading concentrations were calculated as the mass of pDNA per milliliter of undiluted CNBs (μg/NBmL) to capture the ratio of pDNA to the number of CNBs present. To promote even coating, pDNA was first diluted in PBS and placed stirring on 4°C ice with a 7×2 mm magnetic stir bar at ∼250 rpm. CNBs were then added dropwise to the pDNA solution. To allow time for complexation, the pDNA-CNBs mixture was stirred for 15 minutes before use. Plasmid loading was conducted in PBS rather than deionized water to ensure the mixture was compatible with *in vivo* injection. For characterization and loading capacity experiments, loading was conducted at a 100-fold final dilution of CNBs into pDNA, wherein CNBs were first diluted 10-fold in PBS, then added dropwise to the pDNA for a further 10-fold dilution. For cell uptake and transfection experiments, the final dilution factor of CNBs into pDNA varied based on experimental requirements for a desired NB-to-cell ratio.

### Gel electrophoresis of pDNA-CNBs

To assess pDNA loading capacity, CNBs were diluted 100-fold into PBS and loaded at a range of pDNA concentrations (prepared by two-fold serial dilutions, equivalent to 1000, 500, 250, 125, 62.5, 31.25, and 15.6 μg/NBmL. 1000 μg/NBmL is equivalent to 10 μg/mL in total volume) and run on gel electrophoresis to discriminate free pDNA from CNB-complexed pDNA. After pDNA loading, CNBs were mixed with BlueJuice Loading Buffer and loaded into a 1% agarose gel with SYBR Safe DNA Gel Stain. A 1 kb DNA ladder was used as a reference to confirm the size and integrity of the plasmid. Gels were run in 1x Tris-acetate-EDTA buffer (ThermoFisher) for 40 minutes at 120 V, then imaged (Bio-Rad ChemiDoc) with a 0.3 second exposure time.

### Quantification of CNB loading capacity

To quantify the amount of pDNA complexed to the CNBs, a calibration curve correlating absolute pDNA mass to band intensity on gel electrophoresis was generated. A pDNA-only gel with no CNBs was run to represent the 100% un-complexed state (**Fig. S3A**). The pDNA concentrations tested were equivalent to the loading concentrations of the pDNA-CNBs gels (two-fold serial dilutions, beginning at 10 μg/mL, or 1000 μg/NBmL equivalent). After imaging (0.3 second exposure), band intensities of the gel were quantified in ImageJ. Images were converted to 8-bit grayscale, and identical rectangular regions of interest (ROIs) were drawn around each lane. The respective mean grey value versus position profiles were generated using ImageJ’s Plot Profile function (**Fig. S3B**). Data were imported into MATLAB to compute AUC. To separate pDNA band signal from background, a linear boundary was drawn between the curve’s intersection points with standardized position boundaries. Peak area above the boundary was divided by the background area to generate a linear calibration curve relating signal-to-background ratio to pDNA mass (**Fig. S3C**). This calibration curve was then applied to quantify the mass of free pDNA in the pDNA-CNB gels. Given the CNB-complexed pDNA is retained in the sample well, its mass was calculated by subtracting the mass of free pDNA from the total pDNA mass in that condition. Using the average CNB concentration obtained by RMM (2.43×10^11^ NBs/mL), the pDNA mass per individual CNB was obtained.

### Fluorescence microscopy of pDNA on the bubble surface

To visually confirm pDNA loading onto the bubble, the pmax-GFP plasmid was tagged with a Cy5 fluorophore using the Mirus *Label* IT Nucleic acid labeling kit according to manufacturer’s protocol. To accommodate the resolution limits of fluorescence microscopy, larger microbubbles were used as a visual surrogate for the smaller NBs; thus, differential centrifugation was not performed after mechanical agitation of the lipid formulation. Cationic or anionic rhodamine bubbles were loaded with 250 μg/NBmL Cy5-pDNA and placed in a No. 1.5 glass-bottom, 35 mm petri dish (MatTek) covered by a No. 1 glass round coverslip (ThermoFisher Scientific). Bubbles were imaged with a 63x oil objective on the Carl Zeiss microscope to assess colocalization of the rhodamine lipid shell and Cy5 pDNA.

### Agarose phantom production

To house samples for acoustic evaluation *in vitro*, an agarose, tissue-mimicking phantom was created. A phantom mold was designed in CAD software and 3D-printed in heat-resistant acrylonitrile styrene acrylate (ASA) (**Fig. S4A**). The rectangular mold consisted of two interlocking halves for easy removal of the phantom, and a separate insert suspended 3 mm-diameter steel pins to form the sample wells. Multiple wells allowed for simultaneous imaging of three to four separate samples during one ultrasound acquisition. A 1.5% agarose solution in deionized water was heated until fully dissolved, poured into the mold, and cooled at 4°C until fully solidified (**Fig. S4B**). Phantoms were removed from the molds and stored in deionized water at 4°C until use. To prevent cross-contamination across samples, new phantoms were used for each acquisition.

### Ultrasound imaging parameters and signal intensity quantification

All ultrasound imaging, including *in vitro* and *in vivo* experiments, was performed on the Vevo2100 Imaging System (VisualSonics) with a MS-250 transducer (13–24 MHz broadband frequency) at 18 MHz in simultaneous RF nonlinear contrast mode (4% power) and B mode (100% power) at 1 fps. To assess nonlinear contrast intensity, data were exported as AVI files, and contrast in an ROI was measured in ImageJ using the Time Series Analyzer Version 3 plugin. ROIs were kept identical across all *in vitro* samples for comparison and quantified with the Get Total Intensity function. For CNB stability studies, background of PBS-only wells was subtracted. For cell suspension imaging, the cells-only, no CNB control was subtracted. For *in vivo* studies, liver ROIs were drawn for individual mice. Average intensity in the liver ROI was quantified, and pre-injection baseline contrast was subtracted from each frame.

### *In vitro* ultrasound contrast of CNBs

CNB stability under ultrasound was assessed in an agarose phantom at room temperature (RT). The imaging transducer was first immobilized on a stand and horizontally gel-coupled (Aquasonic 100) to the phantom (**Fig. S4C**). To prevent reverberation artifacts, the phantom sat on a silicone acoustic absorber. The centers of the wells were positioned at the focal depth of the transducer to ensure NBs were fully and equivalently subjected to the ultrasound beam. CNBs and 250 µg/NBmL pDNA-CNBs were diluted to the order of 10^9^, 10^8^, or 10^7^ NBs/mL in PBS, then pipetted into respective wells of the phantom. An agarose lid was placed on top to prevent sample evaporation during 500 continuous frames of acquisition. To assess longitudinal stability, undiluted CNBs and rCNBs were place in eppendorf tubes and stored at 4°C at ambient pressure. At various timepoints across 6 days (2, 20, 70, 96, 120, and 144 hours), CNB aliquots were taken from the bulk sample, diluted to 10^9^, 10^8^, or 10^7^ NBs/mL in PBS, and imaged in an agarose phantom.

### Cell culture

PC3 human prostatic adenocarcinoma cells (ATCC) were cultured in complete RPMI1640 medium (Gibco) supplemented with 10% fetal bovine serum (FBS) (Cytiva) and 1% penicillin-streptomycin (P/S) (Gibco) at 37°C, 5% CO_2_. Cells were cultured until 80–90% confluency before passaging and kept below 20 passages. Across all experiments, cells were harvested with 0.25% trypsin/EDTA (Gibco), neutralized with complete RPMI, and pelleted at 300 g for 5 minutes. Mycoplasma testing (InvivoGen MycoStrip) was routinely conducted to ensure cells were pathogen-free before use. Cell counts were obtained manually with a hemocytometer and trypan blue staining.

### Cell uptake kinetics of CNBs

To characterize the kinetics of CNB cellular internalization, PC3 cells were first seeded at 55,000 cells per well in a 24-well plate in complete RPMI + 10% FBS + 1% P/S. After 24 hours, media was removed from the adhered cells, and rCNBs were added at 5,000, 10,000, or 20,000 NBs/cell in incomplete RPMI. Cells were incubated at 37°C until the appropriate endpoint was reached (5, 15, 30, 90, 120, 150, or 180 minutes), then washed twice with PBS to remove any excess NBs. Separate wells were used for each timepoint. Rhodamine signal was imaged by fluorescence microscopy (Zeiss Axio). For pDNA-loaded CNB uptake studies, rCNBs were loaded with 50, 125, or 250 µg/NBmL of pDNA and added to cells at 20,000 NBs/cell.

For flow cytometry, cells were harvested and stained with LIVE/DEAD Fixable Aqua viability dye (ThermoFisher) for 30 minutes at RT. Cells were then washed with PBS and fixed in 4% formaldehyde for 20 minutes at RT. After fixation, cells were washed twice with PBS and resuspended in PBS + 2% FBS. Samples were acquired on the Attune NxT flow cytometer. Rhodamine CNB signal in live cells was quantified using FlowJo version 10.8.1.

### Confocal microscopy of CNB intracellular localization

To visualize intracellular rCNBs, 200,000 PC3 cells were seeded in a 35 mm dish with a 1.5 coverglass bottom (Mattek) and cultured for 48 hours. Media was removed, and 20,000 rCNBs/cell loaded with 50 μg/NBmL of Cy-5-labeled pDNA in PBS were added. CNBs were internalized for 15 minutes at 37°C. Cells were then washed twice with PBS to remove excess NBs. To visualize acidic late-endosomes and lysosomes, samples were stained with 100 nM LysoTracker Yellow HCK-123 (ThermoFisher) in incomplete RPMI for 1 hour at 37°C. Cells were washed with PBS, fixed in 4% formaldehyde for 20 minutes at RT, then stained with DAPI for 30 minutes at RT. The same day, confocal microscopy was conducted on the Leica TCS SP8 gated-STED 3x microscope at 100x magnification. To reduce noise, images were digitally processed in the Huygens Professional deconvolution wizard.

### Ultrasound imaging of PC3 cells after CNB internalization

To confirm internalized CNBs remained echogenic, 150,000 PC3 cells per well were seeded in 12-well plates and cultured for 24 hours in complete RPMI at 37°C. Then, 5,000, 10,000 or 20,000 NBs/cell were added to each well in 400 μL of incomplete RPMI. Cells incubated for either 15 minutes or 1 hour at 37°C while CNBs internalized. After incubation, excess CNBs were removed and cells were washed twice with excess PBS while remaining adherent to the plate. For one group, CNBs were removed after 15 minutes, then cells continued incubating for 1 hour at 37°C in incomplete RPMI to assess signal longevity. For ultrasound imaging, cells were harvested and resuspended in trypsin at 10^6^ cells/mL, and FBS was directly added to a 10% final concentration for neutralization. The cell suspension was then imaged in an agarose phantom.

### *In vitro* therapeutic ultrasound exposure of cell suspensions

Therapeutic ultrasound of PC3 cell suspensions was conducted in 3.4 mL disposable polyethylene transfer pipette (Fisherbrand) submerged in a water bath at RT (**Fig. S6**). After a 15-minute internalization of 20,000 CNBs/cell at 37°C, removal excess CNBs, and 2x washing with PBS, adherent PC3 cells were harvested with trypsin, neutralized with 10% FBS, and resuspended to 100,000 cells/mL in incomplete RPMI. Cells were drawn into the pipette bulb, then placed 2 cm away from a 3 MHz, 1cm^2^ transducer (Sonicator 740, Mettler Electronics) within a water bath. Samples were exposed to therapeutic ultrasound for 0, 10, 20, or 60 seconds at 0.5, 1.0, or 2.2 W/cm^2^ intensity at 20% duty cycle (DC). Nonlinear contrast was evaluated in an agarose phantom immediately after sonication.

### Ultrasound-mediated transfection *in vitro*

To evaluate transfection efficiency *in vitro*, PC3 cells were incubated at 500,000 cells/mL with 20,000 rCNBs/cell pre-loaded with 50 μg/NBmL pmax-GFP pDNA. After pDNA-rCNB internalization for 15 minutes at 37°C, cells were further diluted to 100,000 cells/mL with incomplete RPMI to prevent ultrasound attenuation. To cavitate the CNBs, cells were drawn into the bulb of a plastic pipette, submerged 2 cm away from a 3 MHz, 1cm^2^ transducer (Sonicator 740, Mettler Electronics) in a water bath, and subjected to therapeutic ultrasound for 0, 10, 20, or 60 seconds at 0.5, 1.0, or 2.2 W/cm^2^ intensity at 20% DC. After sonication, cells were plated in 12-well plates and cultured at 37°C. Media was supplemented to 10% FBS 5 hours post-transfection to encourage cell health. After 24 hours to provide time for gene expression, cells were imaged on the Zeiss Axio fluorescent microscope for rCNB and GFP signal. Samples were then harvested for flow cytometry, stained with LIVE/DEAD Fixable Aqua viability dye, fixed in 4% formaldehyde, and read on the Attune NxT flow cytometer. To investigate transfection efficiency with free pDNA and NBs, the equivalent protocol was followed with rhodamine anionic NBs instead of rCNBs at a single ultrasound setting (3 MHz, 20% DC, 2.2 W/cm^2^, 10 seconds).

### CNB contrast kinetics *in vivo*

To assess CNB echogenicity *in vivo*, contrast in mouse livers was recorded after intravenous administration. Male BALB/c mice (Jackson Labs) were first anesthetized with 2% isoflurane and 2 L/min oxygen. To prevent hair interference, abdomens were shaved and treated with hair removal cream (Veet) prior to ultrasound imaging. CNBs were diluted 2-fold in PBS, and 200 μL was directly injected into the tail vein through a 30G needle. CNBs were pre-diluted to mitigate attenuation of the signal. For pDNA-loaded samples, CNBs were first loaded with 50 μL/NBmL of pmax-GFP pDNA at a 2-fold dilution prior to injection. For kinetics studies, the liver was continuously imaged for 50 minutes. To assess contrast after NB cavitation, therapeutic ultrasound was applied 15 minutes after intravenous injection at 3 MHz, 2.2 W/cm^2^, 20% DC for 5 minutes (Sonicator 740, Mettler Electronics). Ultrasound images on the Vevo2100 were recorded immediately before and after therapeutic ultrasound for comparison. All animals were handled according to approved IACUC protocols.

### *In vivo* transfection

Transfection of GFP was assessed in mouse livers. Anesthetized male BALB/c mice received 200 μL of PBS, unloaded CNBs diluted 2-fold in PBS, or 50 μL/NBmL pDNA-CNBs diluted 2-fold in PBS (equivalent to 5 ug pDNA dose per mouse) through bolus tail vein injection. After 15 minutes of circulation time, therapeutic ultrasound over the liver was applied with a 1cm^2^, 3 MHz transducer at 2.2 W/cm^2^ intensity, 20% DC for 5 minutes (Sonicator 740, Mettler Electronics) to cavitate the CNBs. After 24 hours, mice were sacrificed via CO_2_ inhalation, and livers were extracted.

### Anti-GFP staining of liver slices

After *in vivo* transfection, livers were washed with PBS, fixed in 1% formaldehyde for 24 hours at 4°C, soaked in 30% sucrose for 48 hours at 4°C, and frozen in OCT compound (Fisher HealthCare) at −80°C. Next, 8 μm sections of the median liver lobe were cryosectioned (Leica CM3050S) onto glass slides. Excess OCT was washed away with PBS, then slides were permeabilized with 0.1% TritonX-100 (Sigma Aldrich) for 10 minutes at RT, blocked with 10% goat serum in 0.1% Tween20 for 2 hours at RT, and stained with 1:1000 turboGFP rabbit polyclonal primary antibody (OriGene) overnight at 4°C. Excess primary antibody was removed with 5 washes of PBS for 5–10 minutes each, then slides were stained with 1:1000 donkey anti-rabbit AF647 secondary antibody (Abcam) for 1 hour at RT. After 5x PBS washing of excess secondary antibody, slides were stained with 1:1000 DAPI for 15 minutes at RT, then mounted with EverBrite mounting media. Anti-GFP staining with AF647 was conducted instead of looking at raw GFP signal due to high background autofluorescence after fixation. Antibody specificity was confirmed *in vitro* with lipofectamine-transfected PC3 cells (**Fig. S9**).

### Confocal imaging and MFI quantification of histology slides

Stained liver slices were imaged on the Leica TCS SP8 gated-STED 3x confocal microscope at 40x magnification. For each mouse, n = 10 images were acquired across 5-6 sections. Fields-of-view were chosen in the DAPI channel to prevent bias. To evaluate GFP expression, AF647 mean fluorescence intensity (MFI) was quantified in MATLAB and normalized to the cell count in that image. To express the fold increase in GFP expression, MFIs were divided by the average MFI of saline-injected negative controls. CNB rhodamine signal was captured by counting the number rhodamine pixels above threshold, set as 5-times the average rhodamine MFI for that image. Statistical significance computed by one-way ANOVA with multiple comparisons in GraphPad Prism.

## Author contributions

A.A.E., P.N., and L.E.C. conceptualized the project. L.E.C., P.N., and A.A.E. designed experiments. L.E.C. wrote the manuscript with revisions from P.N., A.L., T.B., and A.A.E.. L.E.C., P.N., and M.V.B.J. performed nanobubble characterization experiments. X.H. designed, conducted, and analyzed cryo-EM imaging with laboratory support from M.T.N.. AL. and A.H.K. collected RMM data. A.H.K. and I.B. conceptualized and analyzed GC/MS experiments. L.E.C. performed *in vitro* and *in vivo* transfections. L.E.C. and P.N. performed *in vivo* ultrasound kinetics studies. J.F.E. and M.L.D. advised study design and produced plasmid DNA. L.E.C. analyzed data. M.V.B.J. analyzed DNA loading capacity gels. P.N., S.E.Y., A.L., L.E.C., A.A.E., K.H., F.D., V.F.G., T.B., M.C.P., and R.K.P.B. contributed to nanobubble formulation optimization and *in vivo* study design. All authors reviewed and approved the final manuscript.

## Funding/Acknowledgements

A.A.E. receives support from the National Institutes of Health (NIH) National Institute of Biomedical Imaging and Bioengineering, the National Cancer Institute, and the Department of Defense. L.E.C. received support through the Case Western Reserve University (CWRU) Medical Scientist Training Program (T32 GM007250, T32 GM152319) and the NIH National Center for Advancing Translational Sciences (T32 TR004521). The content is solely the responsibility of the authors and does not necessarily represent the official views of the National Institutes of Health. P.N. acknowledges funding from the 2022 Moderna Global Research Fellowship. X.H. receives research funding from the American Heart Association 2022 Postdoctoral Fellowship (2021AHA000POST000216263) and American Society of Hematology Scholar Award (2024 Basic/Translational Research Fellow). M.T.N. receives research funding from the NIH (HL098217, HL154026). M.L.D., I.B., and J.F.E. receive support from the Cystic Fibrosis Foundation (DRUMM24R0, DRUMM22G0) and the Research Institute for Children’s Health. This work was also supported by the Breakthrough T1D research fund. Confocal microscopy was conducted with the CWRU SOM Light Microscopy Core Facility NIH Grant S10-OD016164. At CWRU, we thank Dr. David Wald’s lab for flow cytometry support and Dr. James Basilion’s lab for cryostat assistance. Figures created with BioRender.com.

