## Supplemental Figures for "Nonviral, ultrasound-triggered gene delivery platform via gas-core cationic nanobubbles"

A

| | | NB formulation | | | | | |
|---|---|---|---|---|---|---|---|
| | | Cationic | | | Anionic | | |
| Lipid | MW (g/mol) | Mass ratio | mM | mol% | Mass ratio | mM | mol% |
| DBPC | 902.36 | 6 | 6.65 | 58.10 | 6 | 6.65 | 58.39 |
| DPPE | 691.96 | 2 | 2.89 | 25.25 | 2 | 2.89 | 25.38 |
| DSPE-mPEG2k | 2790.00 | 1 | 0.36 | 3.13 | 1 | 0.36 | 3.15 |
| DPTAP | 646.50 | 1 | 1.55 | 13.52 | – | – | – |
| DPPA | 670.87 | – | – | – | 1 | 1.49 | 13.09 |

B

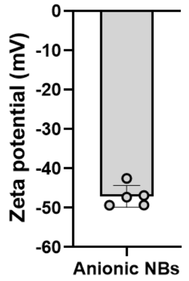


**Figure S1. Cationic and anionic nanobubble formulations. A)** Lipid mass ratios, molarity, and mol percents for the cationic and anionic NB formulations**. B)** Zeta potential of anionic NBs, measured in deionized DEPC-treated water (n = 5).

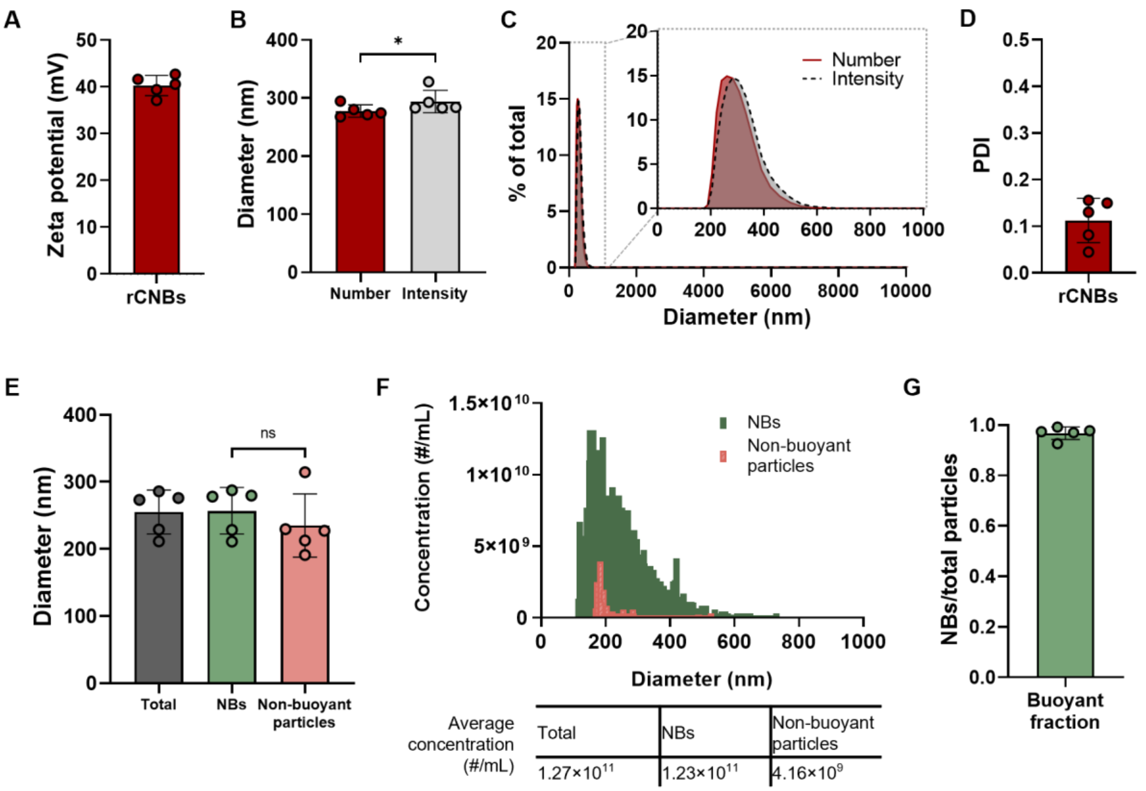


**Figure S2. Characterization of rhodamine-CNBs (rCNBs)**. **A)** Zeta potential (n = 5) in DEPC-treated water. **B)** Peak number-weighted and intensity-weighted diameters by dynamic light scattering (DLS) (n = 5). **C)** Average percent size distribution by DLS with inlay highlighting 0–1000 nm range (n = 5). **D)** Polydispersity index (PDI) by DLS (n = 5). **E)** Diameter of buoyant NBs, non-buoyant particles, and total particles by resonant mass measurement (RMM) (n = 5). **F)** Concentration vs. diameter distribution of NBs and non-buoyant particles by RMM (n = 5). **G)** Fraction of buoyant NBs to total particles by RMM (n = 5).

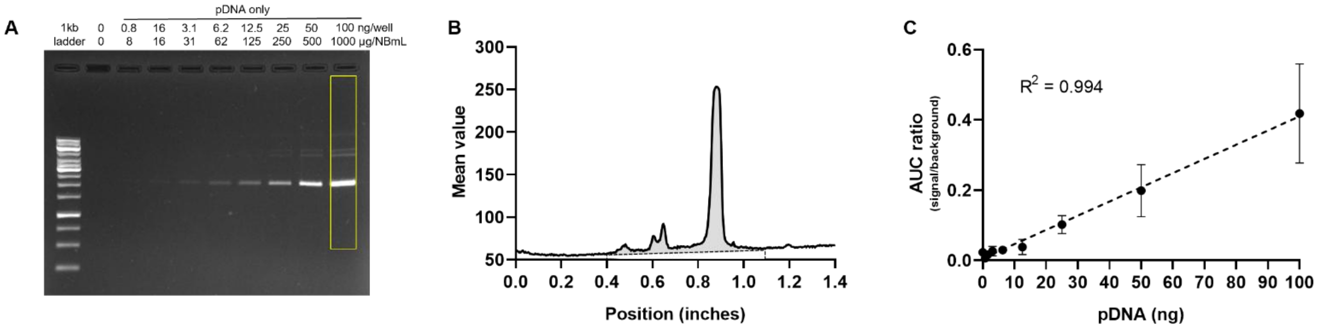


**Figure S3. Gel electrophoresis of pDNA without CNBs and calibration curve for loading capacity quantification. A)** Agarose gel of pDNA without CNBs at equivalent concentrations to those in Fig. 3A (0–1000 µg/NBmL). **B)** Representative mean value vs. position plot in the gel region of interest (ROI). Area under the curve (AUC) of the pDNA peak shaded. **C)** Calibration curve of pDNA concentration vs. AUC signal-to-background ratio in the pDNA-only gel ROIs, $R^2$ = 0.994 (n = 3).

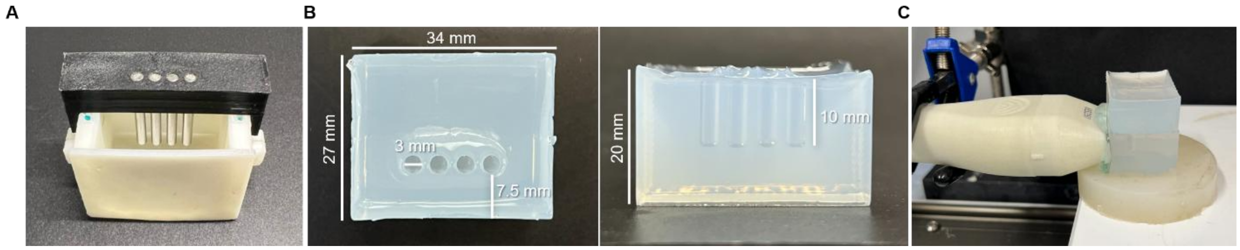


**Figure S4. Agarose phantom and *in vitro* ultrasound imaging setup. A)** Phantom mold, 3D printed in acrylonitrile styrene acrylate (ASA) and fitted with an insert holding 3 mm diameter steel pegs. **B)** Agarose phantom with dimensions, top and side views. **C)** Ultrasound imaging acquisition set-up, with transducer horizontally gel-coupled to the phantom. Agarose lid placed on top to prevent sample evaporation during acquisition.

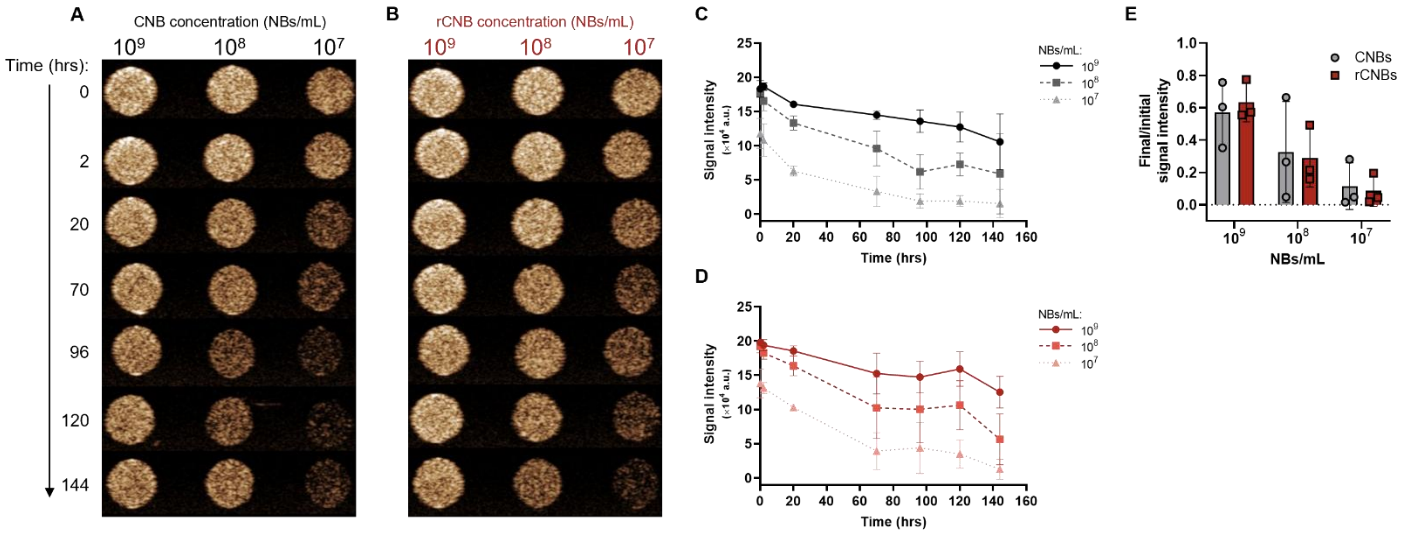


**Figure S5. Ultrasound contrast longevity of CNBs and rhodamine-CNBs (rCNBs).** Representative nonlinear contrast images and signal intensity quantification (n = 3) of **A,C)** CNBs and **B,D)** rCNBs over time. **E)** Signal intensity at t = 144 hours divided by signal intensity at t = 0 hrs. Undiluted samples were stored at 4°C, and aliquots were extracted and diluted to the order of $10^9$, $10^8$, or $10^7$ NBs/mL in PBS at each timepoint before imaging. PBS background subtracted.

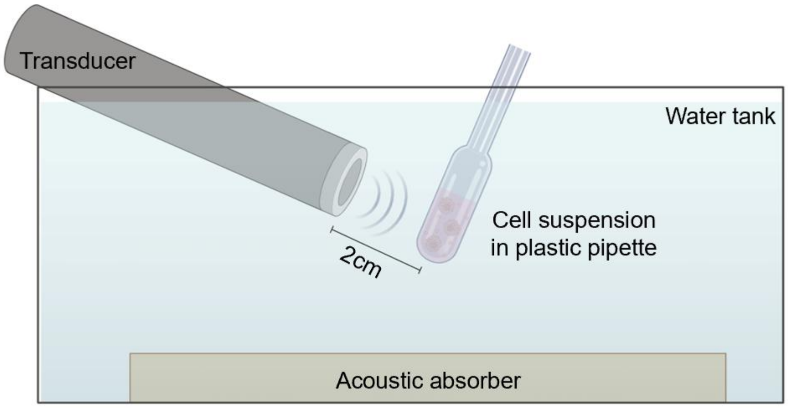


**Figure S6*. In vitro* therapeutic ultrasound setup.** Cell suspensions were drawn into the bulb of a plastic pipette and sonicated in a water bath from a 2 cm distance. A silicone acoustic absorber lined the bottom of the tank to mitigate reflections.

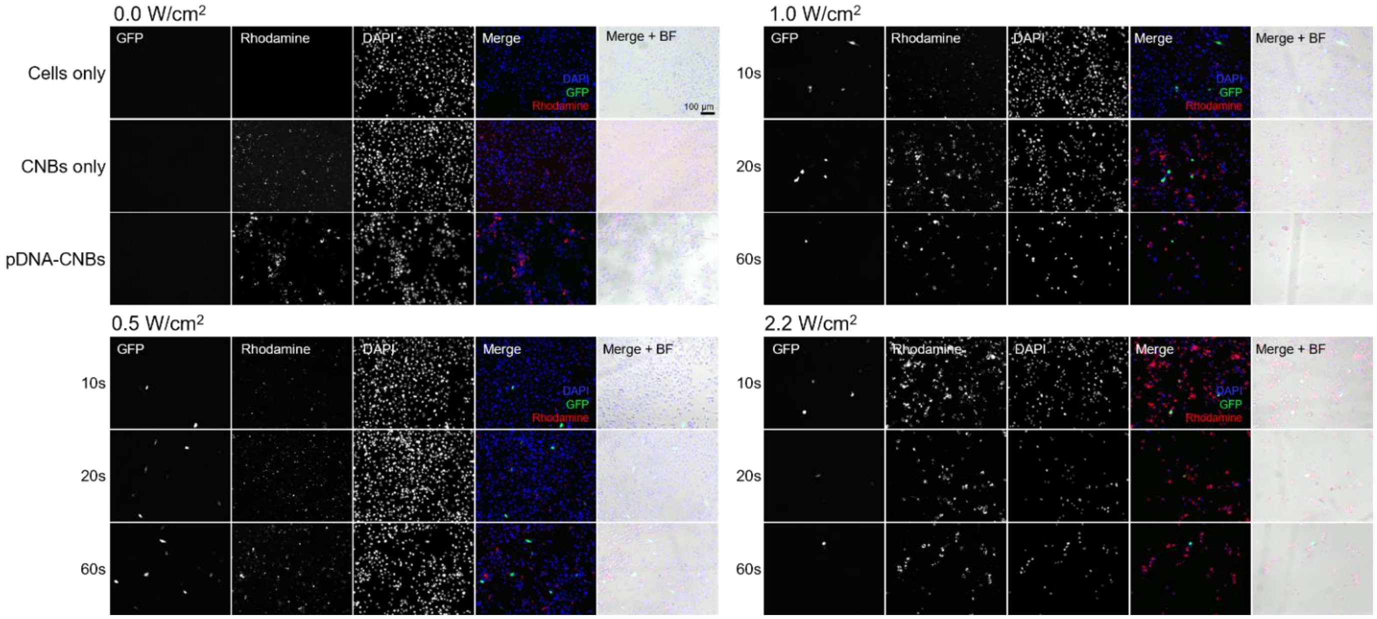


**Figure S7. Fluorescence microscopy images of PC3 cells 24 hours post *in vitro* transfection**. Cells internalized rhodamine-CNBs loaded with pmax-GFP pDNA, and 3 MHz therapeutic ultrasound was applied at varying durations (10, 20, or 60 seconds) and intensities (0.5 W/cm$^2$, 1.0 W/cm$^2$, or 2.2 W/cm$^2$) at 20% duty cycle.

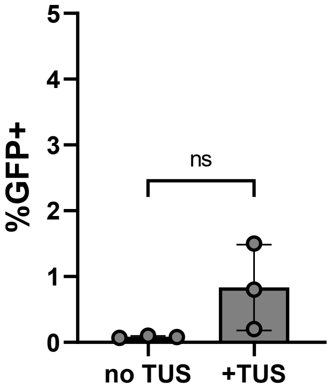


**Figure S8.** ***In vitro* transfection in PC3 cells with anionic NBs**. Percent GFP positive cells, 24 hours post *in vitro* transfection with 50 µg/NBmL pmax-GFP pDNA and anionic NBs, with and without therapeutic ultrasound (TUS) at 3 MHz, 2.2 W/cm$^2$, and 20% duty cycle for 10 seconds (n = 3).

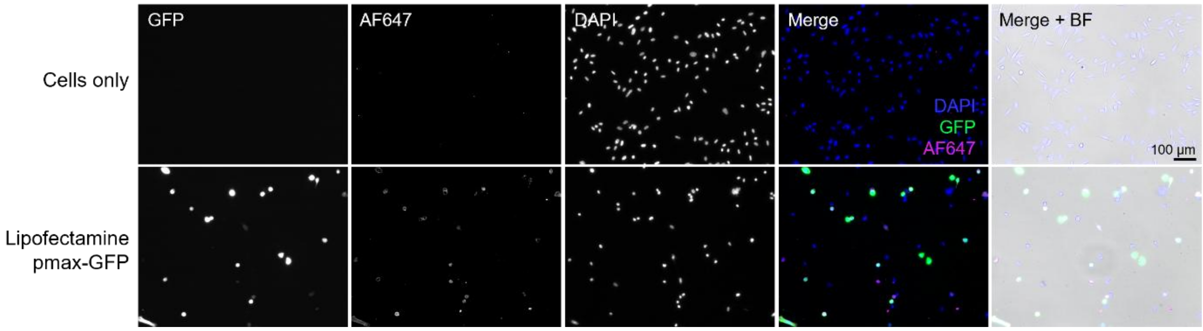


**Figure S9. Anti-GFP antibody staining of PC3 cells *in vitro*.** To confirm antibody specificity for GFP, PC3 cells were transfected with lipofectamine and pmax-GFP pDNA according to manufacturer's protocol. Transfected cells and an untransfected, cells-only control were then stained with anti-GFP 1° and AF647 2° antibodies. Scale bar 100 µm.
